# Fine-tuning the Spike: Role of the nature and topology of the glycan shield in the structure and dynamics of the SARS-CoV-2 S

**DOI:** 10.1101/2021.04.01.438036

**Authors:** Aoife M. Harbison, Carl A. Fogarty, Toan K. Phung, Akash Satheesan, Benjamin L. Schulz, Elisa Fadda

## Abstract

The dense glycan shield is an essential feature of the SARS-CoV-2 spike (S) architecture, key to immune evasion and to the activation of the prefusion conformation. Recent studies indicate that the occupancy and structures of the SARS-CoV-2 S glycans depend not only on the nature of the host cell, but also on the structural stability of the trimer; a point that raises important questions about the relative competence of different glycoforms. Moreover, the functional role of the glycan shield in the SARS-CoV-2 pathogenesis suggests that the evolution of the sites of glycosylation is potentially intertwined with the evolution of the protein sequence to affect optimal activity. Our results from multi-microsecond molecular dynamics simulations indicate that the type of glycosylation at N234, N165 and N343 greatly affects the stability of the receptor binding domain (RBD) open conformation, and thus its exposure and accessibility. Furthermore, our results suggest that the loss of glycosylation at N370, a newly acquired modification in the SARS-CoV-2 S glycan shield’s topology, may have contributed to increase the SARS-CoV-2 infectivity as we find that N-glycosylation at N370 stabilizes the closed RBD conformation by binding a specific cleft on the RBD surface. We discuss how the absence of the N370 glycan in the SARS-CoV-2 S frees the RBD glycan binding cleft, which becomes available to bind cell-surface glycans, potentially increases host cell surface localization.

## Introduction

Spike (S) glycoproteins mediate the adhesion and fusion of enveloped viruses to the host cell, initiating viral infection^1–3^. The interaction with the cell-bound receptor leads to a complex conformational change, dependent on the S architecture^4, 5^, that terminates with the fusion of the viral envelope with the host cell’s membrane, giving the virus access to the cellular machinery for replication^3^. Viral envelope S are heavily coated with a dense layer of complex carbohydrates, also known as a glycan shield, that performs different intrinsic and extrinsic functions, from modulating protein folding, stability and trafficking, to masking the virus from the immune system and mediating contacts with lectins and antibodies^1^. The severe acute respiratory syndrome coronavirus 2 (SARS-CoV-2) has a trimeric S glycoprotein^6, 7^ protruding from the viral envelope surface^8, 9^. The SARS-CoV-2 S has 22 N-glycosylation sequons per protomer, of which at least 18 appear to be consistently occupied in different constructs^8, 10–15^, and O-glycosylation sites with significantly lower occupancy^11, 12, 16^.

A unique feature of the SARS-CoV-2 S glycoprotein’s architecture is the key role of the glycan shield in its activation mechanism^17, 18^. Binding of SARS-CoV-2 S to its primary receptor, namely the angiotensin-converting enzyme 2 (ACE2)^19, 20^, requires the opening of one (or more) receptor binding domains (RBDs), which need to emerge from the glycan shield to become accessible^9, 17, 18, 21–23^. The RBD opening creates a cavity within the SARS-CoV-2 S prefusion trimer’s structure^6, 7^, see **Figure 1**, which extends deeply into the trimer’s core. In the absence of strategically positioned N-glycans as a support^17^, upon RBD opening this large pocket would be filled by water molecules, likely weakening the S prefusion structural integrity, especially considering the SARS-CoV-2 S pre-cleaved polybasic furin site at the S1/S2 boundary^24, 25^. Multi-microsecond molecular dynamics (MD) simulations supported by biolayer interferometry experiments^17^ have shown that this structural weakness is effectively recovered by the N-glycan at position N234, where a site-specific large oligomannose^8, 10, 14, 15^ is able to fill the cavity, supporting the RBD open conformation^17^. Furthermore, MD simulations have also shown that the N-glycans at positions N165 and N343, see **Figure 1**, are directly involved in important interactions with residues of the open RBD, supporting its open conformation^17^ and mediating (or gating) its transition from open to closed^18^, respectively.

**Figure 1.**
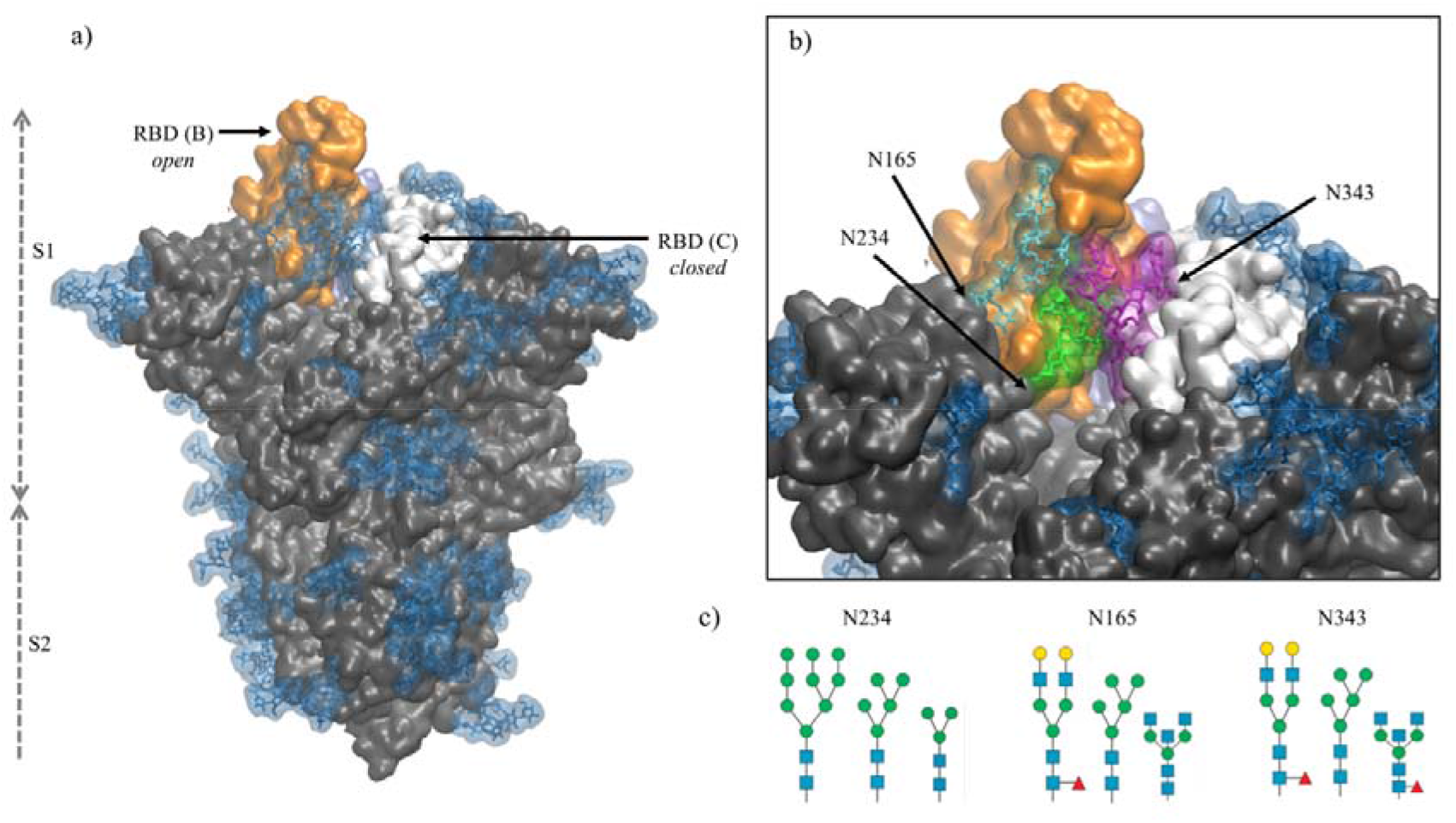
**Panel a)** Structure of the fully glycosylated SARS-CoV-2 S (PDBid 6VYB) ectodomain^*7*^. The protein is shown in grey with the RBDs of chain B and C highlighted in orange and white, respectively; the glycan shield is highlighted in blue. **Panel b)** Close up on the open pockets with the N-glycans at the strategic positions N234, N165, and N343 highlighted in green, cyan and purple, respectively. **Panel c)** N-glycans considered in the different models studied in this work, represented through the SNFG^*26*^ and drawn with DrawGlycan^*27*^ (http://www.virtualglycome.org/DrawGlycan/). Molecular rendering done with VMD^*28*^.

The crucial role of the N-glycans at N234, N165 and N343 is exerted through the contacts these structures can make with protein residues, both in the open RBD (of chain B, following the PDBid 6VYB nomenclature) and in the adjacent closed RBD (chain C) at either side of the empty cleft^17^, see **Figure 1**. Their ability to engage in effective interactions is intrinsically linked to the nature, size, sequence and branching of the N-glycans at these sites, opening the floor to a broader discussion on the relative structural stability and dynamics of different S glycoforms. This is a very important point to explore, especially in view of the design of specific antiviral therapeutic strategies targeting glycosylation^29^, yet a very difficult (if not impossible) one to systematically address experimentally.

In this work, we present the results of a set of multi-microsecond MD simulations of different SARS-CoV-2 S glycoforms aimed at characterizing the effect of changes in the type of glycosylation at positions N234, N165 and N343, see **Figure 1**, while the rest of the glycan shield is represented consistently with a stable recombinant S prefusion trimer (S_trimer_)^10, 17^. More specifically, we investigated if shorter oligomannoses structures at N234, such as Man5 and paucimannose (Man3), see **Figure 1**, could occupy the empty cleft in the open SARS-CoV-2 S as effectively as larger oligomannoses such as Man9, characteristic of a highly stable prefusion S_trimer_^10^ As an important note, Man5, rather than Man7/8/9 is found to be present, or even to be the dominant glycosylation type at N234 in the virus, vaccine epitopes and in some recombinant S constructs^8, 11, 30^. We also explored how the role of oligomannose structures at N234 is supplemented by the complex N-glycans at N165 and N343, which can form a tight network of glycan-glycan and glycan-protein interactions that stabilize the orientation and dynamics of the open RBD across different possible orientations.

Finally, we also investigated the effect of a mutation, unique to the SARS-CoV-2 Wuhan-Hu-1 (NCBI Reference Sequence: NC_045512.2) and derived strains^31^, that changes the RBD glycan shield’s topology. More specifically, in SARS-CoV and MERS^32^, as well as in the bat RaTG13 and pangolin CoV SARS-CoV-2 variants^25, 33, 34^, position N370 on the RBD is part of an occupied NST sequon^32^, which is lost in SARS-CoV-2 due to a T372A mutation. We performed ancestral sequence reconstruction of selected SARS S sequences to investigate the gain and loss of glycosylation sequons during S evolution. To address the effect of this change in the glycan shield’s topology, we re-introduced the N-glycan at N370 in the SARS-CoV-2 S native sequence and ran MD simulations to study its effect on the stability of both, the open and closed RBDs. Our simulations indicate that within the SARS-CoV-2 S architecture, glycosylation at N370 does not interfere with the glycan network at N234, N165 and N343, but actively contributes to it by stabilizing very effectively the RBD open conformation.

Interestingly, analysis of the closed protomers shows that the N370 glycan from RBD (A) is firmly bound to the RBD (C) surface, where it occupies a specific cleft. This interaction results in tying the closed RBD (C) to the adjacent closed RBD (A), very much like the laces in a shoe, thus potentially hindering the opening of the RBDs. Based on these findings, we propose that the recent loss of N-glycosylation at N370 allows for a higher availability of open S conformations by lowering the energetic cost of the opening reaction, which is likely to be beneficial by providing higher infectivity of SARS-CoV-2 relative to closely related variants carrying this sequon. We also discuss how the cleft on the closed RBD surface in SARS-CoV-2 S, which is occupied by the N370 glycan in S glycoforms with the sequon, may be used by other glycans found on the cell surface, such as glycosaminoglycans^35–40^, sialogangliosides, and blood group antigens^39^, where these interactions may contribute to increasing the S cell-surface localization.

## Results

In this section we will present the results of extensive conformational sampling based on multi-microsecond MD simulations of SARS-CoV-2 S models with different glycosylation at N234, N165 and N343, see **Table 1**. Specifically, we determined the effects of site-specific glycan structure on the SARS-CoV-2 S ectodomain’s structure and dynamics by systematically reducing the size of the oligomannose at N234; from Man9 (N234-Man9), found in highly stable prefusion SARS-CoV-2 S constructs^10, 12, 30^, and also studied in previous work^17, 41^, we progressed to a shorter Man5 (N234-Man5), found on the virus and on other SARS-CoV-2 S constructs^11, 30^, and to paucimannose (N234-Man3) as a hypothetical size limit. In these models the glycosylation at N165 and N343 is biantennary complex (FA2G2), and in one case we have also considered bisecting GlcNAc F/A2B forms at N343 and N165, respectively, see **Figure 1 and Table S.1**. The results obtained for the N234-Man5 models were compared to a SARS-CoV-2 S model with a uniform immature glycosylation^42^, i.e. in which all N-glycans are Man5 (all-Man5). Finally, to assess the effect of loss of glycosylation at N370 on SARS-CoV-2 S, we added a complex N-glycan (FA2G2) at N370 in the N234-Man9 model with bisecting GlcNAc (F/A2B) N-glycans at N165 and N343, respectively. In all of these models the glycosylation at all sites other than the ones listed above is consistent and the same as the profile experimentally determined for the stable prefusion S_trimer_^10^, see **Table S.1**. Results are based on the analysis of multiple uncorrelated MD trajectories (replicas) run in parallel for each model (see details in the Computational Methods section in Supplementary Material).

**Table 1.** Orientation of the RBD in different SARS-CoV-2 S glycoforms in terms of average lateral and axial angle values relative to the SARS-CoV-2 S ectodomain, as described in ref.^*17*^. Standard deviation values are indicated in parentheses. Total simulation time for each replica is indicated in parentheses. Note: the first 300 ns of each trajectory were considered as part of conformational equilibration and were omitted from the analysis.

| Model System | Replicas | Lateral Angle (°) | Axial Angle (°) |
| --- | --- | --- | --- |
| N234-Man5 | R1 (2 μs) | 0.0 (2.4) | 2.9 (1.1) |
| | R2 (2 μs) | -6.7 (2.9) | 4.8 (0.9) |
| | R3 (1.5 μs) | 2.7 (3.7) | 5.9 (0.9) |
| N234-Man3 | R1 (2 μs) | 4.4 (3.1) | 1.8 (1.1) |
| | R2 (1.9 μs) | 1.3 (2.5) | -0.1 (1.0) |
| | R3 (1.4 μs) | 2.5 (2.5) | 6.7 (1.1) |
| N234-Man9 | R1 (2 μs) | 3.4 (2.0) | -0.5 (0.7) |
| | R2 (1.4 μs) | -0.7 (2.0) | 4.8 (0.7) |
| N234-Man9 (+ N370) | R1 (1.4 μs) | -3.4 (2.4) | 6.0 (0.7) |
| | R2 (1.3 μs) | -0.6 (2.1) | -1.0 (0.7) |
| All-Man5 | R1 (1.4 μs) | -14.1 (2.6) | 3.2 (0.9) |
| | R2 (1.1 μs) | -3.5 (3.6) | 6.9 (1.0) |
| | R3 (1.9 μs) | -1.9 (6.4) | -0.2 (3.8) |
| N234-Man9* (Man5 N165/N343) | R1 (2.1 μs) | 0.2 (2.5) | 7.3 (1.0) |

### N234-Man5

The structure and dynamics of the RBD (residues 330 to 530) from a model with a Man5 at N234 and FA2G2 at N165 and N343 was analysed based on three replicas with sampling times indicated in **Table 1**. Note that the first 300 ns of each replica were omitted from the analysis to allow conformational equilibration, see **Figure S.1**. The dynamics of the RBD is rather complex and for consistency with previous work^17^ we defined it here in terms of lateral and axial angle values populations. The lateral angle indicates an in-plane motion of the open RBD along a hypothetical circle centred on the central helices of the spike. This angle is defined by three points, corresponding to 1) the centre of mass of the open RBD core β-sheets at frame 0, i.e. the orientation of the RBD in PDB 6VYB, 2) the centre of mass of the top section of the central helices (CH), and 3) the centre of mass of the open RBD core β-sheets at each frame along the trajectory^17^. The axial-angle defines a tilting motion of the open RBD either away from (negative values) or toward (positive values) the central helices of the spike. The axial angle is defined by 1) the centre of mass of the open RBD core β-sheets, 2) the centre of mass of the central helices, and 3) the centre of mass of the top section of the central helices^17^. A graphical representation of the axial and lateral angle coordinate frame is shown in Figure 2.

**Figure 2.**
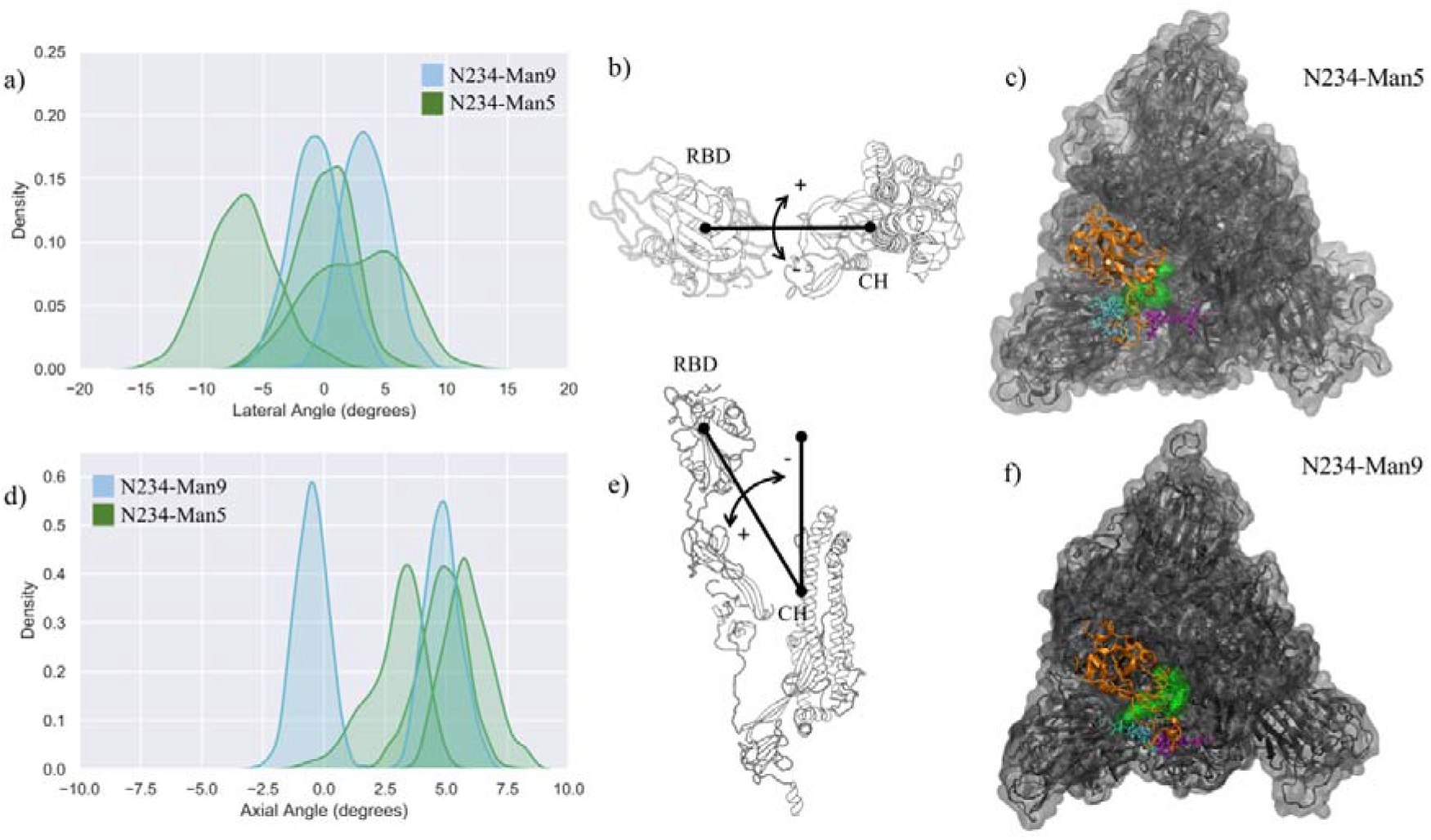
**Panels a)** and **d)** Kernel density estimation (KDE) analysis of the lateral and axial angles distributions calculated through the uncorrelated MD trajectories obtained for N234-Man5 (green), replicas R1-3, and for N234-Man9 (cyan), replicas R1,2 for comparison. **Panels b)** and **e)** Graphical representation of the lateral (b) and axial (e) angles. Displacements relative to the initial trajectory frame are measured in terms of positive and negative values. The centre of mass of the RBD and of the central helices (CH) are used as reference points. **Panels e)** and **f)** Top view on an equilibrated snapshot of the N234-Man5 (R3) S and N234-Man9 (R2) S, respectively. Man5/9 at N234 are shown in green, while the N-glycans at N165 and N343 are shown in cyan and purple, respectively. All other glycans are not represented for clarity. The RBD of chain B is shown in orange, while the rest of the protein is in grey. Data analysis and graphs were done with seaborn (www.seaborn.pydata.org) and molecular rendering with pymol (www.pymol.org) and VMD^*28*^.

Results shown in **Figure 2** and **Table 1** indicate that in the N234-Man5 S model the open RBD can access a larger conformational space relative to the N243-Man9 model along the lateral displacement coordinate, while it is still able to adopt a stable open conformation, indicated by the axial tilt values. To note, the results from the MD simulations of the N243-Man9 ectodomain model presented here agree with the results obtained for the whole N243-Man9 S glycoprotein model^17^. The N234-Man5 S open conformations obtained from the conformational sampling scheme we used are slightly diverse, encompassing different degrees of opening, as shown in **Figures S.1** and **S.2**. More specifically, the RBD can be ‘wide-open’, see in **Figure 2 panel b**, i.e. in a similar orientation seen in the cryo-EM structures^6^, stabilized by interactions involving different RBD residues and the N165 and N234 N-glycans, while the N343 N-glycan is relatively free on the opposite side of the pocket, or alternatively the RBD can be in an intermediate conformation between the wide-open and the closed. In the latter the N-glycans at all three positions N165, N234 and N343 are involved in complex interactions with protein residues in the RBD, namely with the loop from L460 and F489, and in and around the pocket, and with each other, see **Figure S.3**. Within this framework, the higher lateral degree of flexibility of the open RBD in the N234-Man5 S model is due to the smaller size of the Man5 glycan and its specific conformational propensity relative to Man9, which adopts a more ‘tree-like’ conformation, supported by extensive inter-arm interactions^43^, that fills the pocket much more effectively, see **Figure 2 panels e** and **f**. More specifically, the Man5 at N234 appears to be less competent than Man9 at forming interactions that bridge both sides of the pocket as it accesses the cavity left open by the RBD, while it mainly interacts with residues at the base of the open RBD, leaving it ‘unhinged’ when in the “wide-open” conformation, relative to the N234-Man9 model. As an interesting point to note, in all the trajectories the core fucose of the N343 FA2G2 glycan is exposed to the solvent, potentially allowing for its recognition. This is in agreement with cryo-EM studies reporting interactions involving the core fucose of the N-glycan at N343 and human neutralizing antibodies^44^.

To further assess the role of the type of glycosylation around the N234-Man5 in the RBD dynamics, we ran three MD simulations of a model with uniform Man5 glycosylation (all Man5 model), see **Table 1** and **Figure S.4**. The results indicate that the replacement of the complex glycans with immature Man5 structures is problematic for the stability of the open RBD. More specifically, the shorter size and 3D architecture of the Man5 at N343 does not allow it to engage effectively with residues in the RBD, while the Man5 at N234, as seen for the N234-Man5 model, is not able to effectively engage with the closed RBD (chain C) that flank the opposite side of the empty pocket. These results agree with recent work indicating that immature glycosylation, achieved in in GnTI−/− mutant cells, leads to a less competent S glycoprotein relative to the fully glycosylated variant^42^. For comparison we tested a N234-Man9 model where only the N-glycans at N165 and N343 were Man5. The results shown in **Table 1** and **Figure S.5** indicate that the presence of a large oligomannose structure at N243 helps recover the instability by more effectively occupying the empty pocket. In the equilibrium conformation sampled through 2.1 μs, the RBD conformation is more closed, shown also by the positive value of the axial angle, see **Table 1**, where both Man5 at N165 and N343 can interact with each other and with the RBD residues, see **Figure S.5**.

### N234-Man3

To gauge the implications of the size of the oligomannose at N243 for the orientation and dynamics of the open RBD, we studied a model were N243 is modified with a paucimannose (Man3). Note, to our knowledge, the presence of Man3 at this or at any sequon in the SARS-CoV-2 S has not been detected to date^8, 10–13, 30, 45, 46^; nevertheless, as Man3 is one of the smallest oligomannose structures, this S glycoform represents a case study to account for the effect of the reduction in the size of the N-glycan to an extreme at this strategic position. The results shown in **Table 1** and **Figure 3** in addition to the visual analysis of all trajectories, show that Man3 at N234 is less competent than Man 5 and Man9 in supporting a fully wide-open RBD. In only one of the three replicas, namely R3 shown in **Figure 3 panels b** and **d**, the Man3 accesses the interior of the open pocket, although its size does not allow for it to form any interactions that contribute to stability. In both R1 and R2, Man3 interacts with residues outside or at the edge of the pocket, as shown in **Figure S.6**.

**Figure 3.**
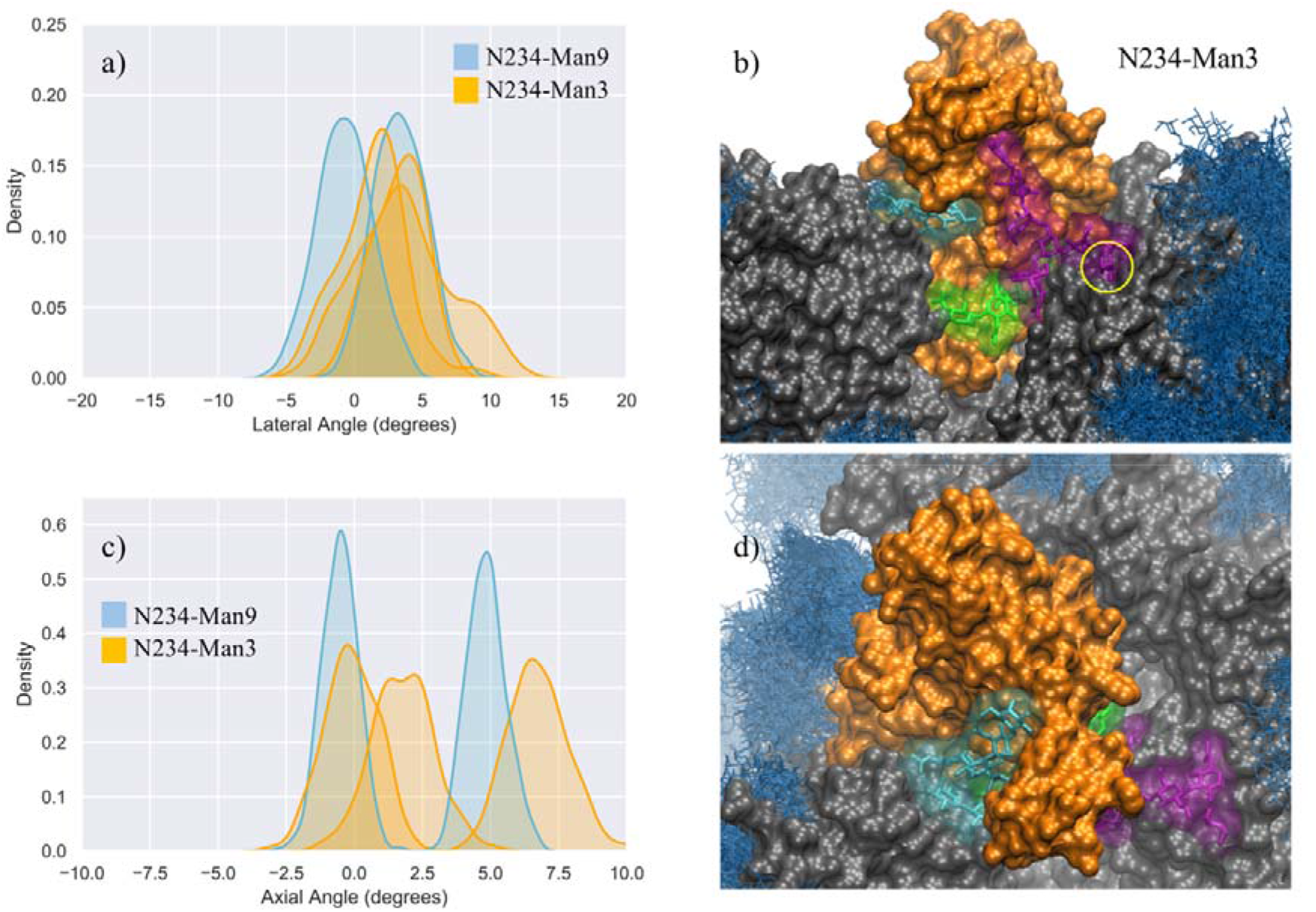
**Panels a)** and **c)** Kernel density estimation (KDE) analysis of the lateral and axial angle distributions calculated through the uncorrelated MD trajectories obtained for N234-Man3 (orange), replicas R1-3, and for N234-Man9 (cyan), replicas R1,2 for comparison. **Panels b)** and **d)** Close-ups on a representative snapshot of the N234-Man3 (R3) simulation from the side and top, respectively. Man3 at N234 is shown in green, while the N-glycans at N165 and N343 are shown in cyan and purple, respectively. The solvent accessible surface of the open RBD is shown in orange, while the rest of the protein is shown in grey. All other glycans are shown in blue, as an overlay of snapshots collected every 10 frames. The position of the core fucose in the FA2G2 N-glycans at N343 is highlighted within a yellow circle. Data analysis and graphs were done with seaborn (www.seaborn.pydata.org) and molecular rendering with VMD^*28*^.

In the N234-Man3 model, the open RBD does not show any significant lateral excursions, see **Figure 3 panel a**, as its position is firmly held in place by interactions with the complex N-glycans at N165 and N343, which contribute to pulling it towards a conformation intermediate near to a closed RBD, shown in **Figure S.2** and **S.6**. In this conformation the N165 and N343 N-glycans interact extensively with residues in the receptor binding motif (RBM), possibly preluding to a final closing, as reported in earlier work, showing the gating activity of the glycans at N343^18^. When the RBD is near closing there is very a very limited degree of freedom in the lateral angle coordinate. As an interesting point, when the FA2G2 N-glycan at N343 interacts with the RBD, its core fucose becomes less accessible to a potential recognition, see **Figure 3 panel b**.

### N234-Man9 with FA2G2 at N370

Analysis of the glycan shield topology through the ancestral sequence reconstruction of select SARS S sequences, shown in **Figures 4 panel d**, indicates the loss of a sequon at N370 in the SARS-CoV-2 Wuhan-Hu-1 strain, due to a mutation to NSA of the conserved NST present in closely related strains of current interest, namely SARS-CoV, the bat RaTG13G S and the pangolin CoV S. To study the effect on SARS-CoV-2 S structure and dynamics of this additional ancestral glycan at N370, we restored the sequon at N370 and glycosylated this position with a complex FA2G2 N-glycan, consistently with data reported in the literature for SARS-CoV^47^. In this model, position N234 is modified with Man9, while positions N343 and N165 are glycosylated with complex N-glycans with a bisecting GlcNAc and core fucosylation at N343 (FA2B), and without core-fucosylation at N165 (A2B). Our results, shown in **Figure 4** and **Table 1**, indicate that the wide-open RBD conformation in the N370-glycosylated SARS-CoV-2 S is as stable as the one observed for the corresponding N234-Man9, where the N-glycan at N370 fills the interior of the empty pocket together with the Man9. In this model, the Man9 at N234 is only able to access the entrance of the pocket due to steric hindrance with the N370 glycan that occupies the core. This suggest that once the RBD opens, an S glycoform with glycosylated N370 would be highly competent in exposing the RBD to the ACE2 receptor, as the SARS-CoV-2 S with a large oligomannose at N234.

**Figure 4.**
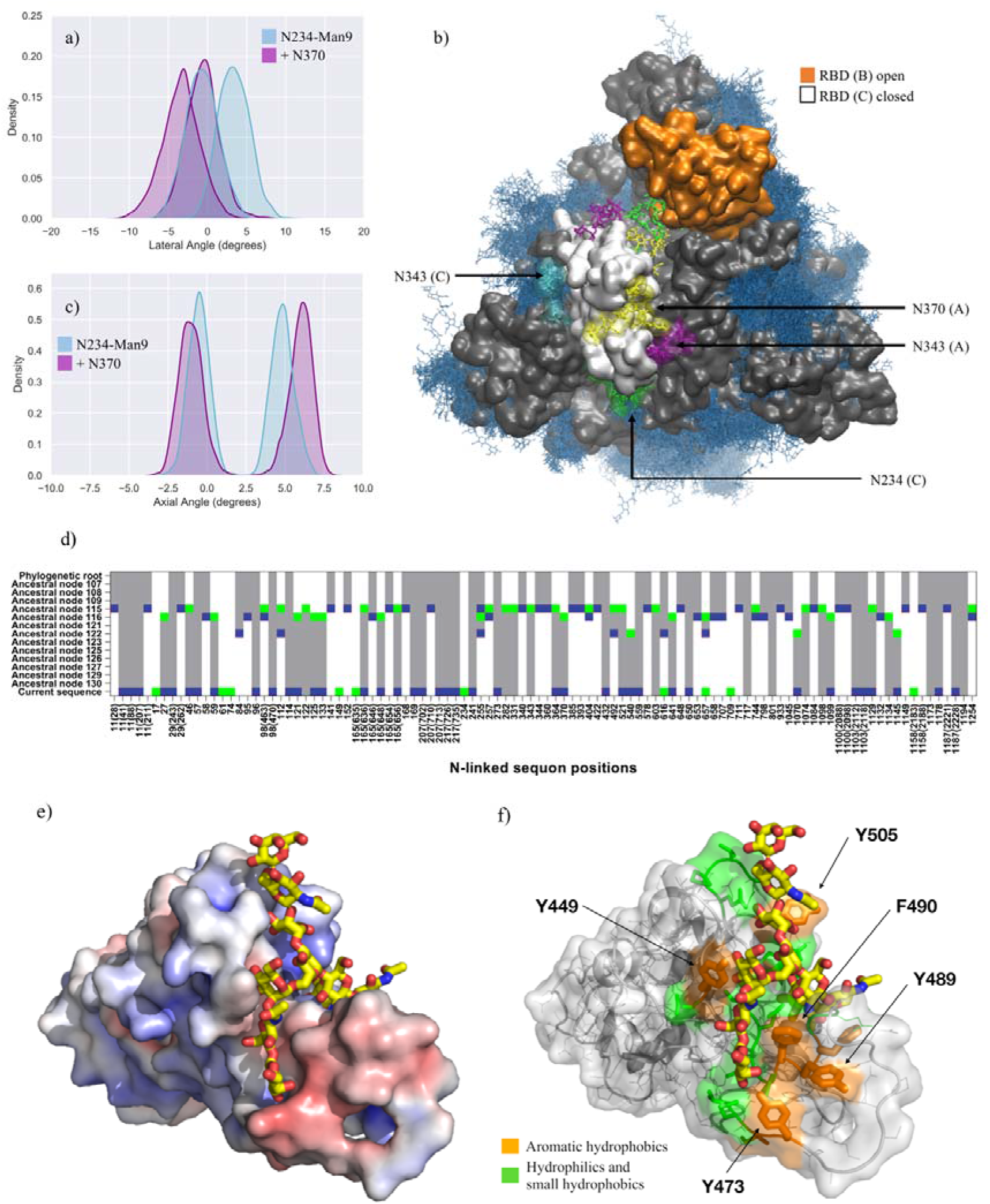
**Panels a)** and **c)** Kernel density estimation (KDE) analysis of the lateral and axial angle distributions calculated through the uncorrelated MD trajectories obtained for N370-glycosylated N234-Man9 (purple), replicas R1,2, and for N234-Man9 (cyan), replicas R1,2 for comparison. **Panel b)** Close-ups on a representative snapshot of the N370-glycosylated N234-Man3 (R2) simulation from top. Man9 at N234 (B,C) is shown in green, while the complex N-glycans at N165 (B,C), N343 (C,A) and N370 (C,A) are shown in cyan, purple and yellow, respectively. The surface of the open RBD (B) is shown in orange and of the closed RBD (C) is shown in white; the rest of the protein is shown in grey. All other glycans are shown in blue, as an overlay of snapshots collected every 10 frames. Data analysis and graphs were done with seaborn (www.seaborn.pydata.org) and molecular rendering with VMD28. **Panel d)** Gain, loss, and retention of N-glycosylation sequons through the evolution of SARS-CoV-2 S as inferred from ancestral sequence reconstruction based on selected coronavirus spike proteins. The predicted ancestral sequence is at the top and the current SARS-CoV-2 S sequence at the base. Colours show gain (green), retention (grey), or loss (blue) of a sequon at a specific amino acid position at each phylogenetic node. N-glycosylation sequon positions are numbered as in the current SARS-CoV-2 sequence. Positions in the current SARS-CoV-2 sequence that are aligned with multiple sequons in reconstructed ancestral sequences due to insertion/deletion events also include their position in the multiple sequence alignment in parentheses. **Panel e)** Close-up of the RBD C bound to the N370 FA2G2 N-glycan represented with sticks and yellow C atoms. The RBD C is represented through a solvent accessible surface colorised based on the electrostatic potential calculated with the APBS plugin in pyMol (www.pymol.org). Darker shades of blue indicate increasingly positive charge, white indicate neutral charge and increasingly red shades indicate negatively charged regions. **Panel f)** Representation of the RBD C bound to the N370 FA2G2 N-glycan in the same orientation as in Panel e) highlighting potentially critical residues for binding. Molecular rendering done with pyMol.

The analysis of the dynamics of the closed RBDs of protomers A and C (PDB 6VYB numbering) shows that the N370 N-glycan of RBD (A) is tightly bound to the surface of the adjacent closed RBD (C), threading the two closed RBDs together, see **Figure 4 panel b, e** and **f** and **Figure S.7**. The 3D architecture of the complex FA2G2 N-glycan at N370, characterized by independent dynamics of the arms^48^, allows for stable interactions of the (1-6) arm within a cleft in the RBD, flanked by residues between N448 and Y453 on one side and F490 and Y495 on the other, see **Figure 4 panel e** and **f** and **Figure S.7**, that support binding through hydrogen bonding and hydrophobic interactions with different monosaccharide units along the (1-6) arm. Because of the stability of the N370 glycan-protein interaction in the closed RBD, N370 glycosylation may hinder the opening mechanism. Thus, the loss of glycosylation at N370 likely contributes to enhancing the binding activity of S and infectivity of SARS-CoV-2 relative to other variants with an N-glycosylation sequon at this position.

## Discussion

The SARS-CoV-2 S glycosylation profile with expression in (or infection of) mammalian cells has been reported by several studies^8, 10–15, 45^, almost all of which find a large oligomannose N-glycan, such as Man9-7, as the most common structure at N234. This is especially true in highly stable prefusion SARS-CoV-2 S trimer glycoforms^10, 30^, which bear the 2P mutation^6, 49^. Meanwhile, a shorter Man5 appears to be present or even the dominant structure at N234 in the virus^8, 30^ and in the secreted ChAdOx1 nCoV-19 (AZD1222) vaccine epitope^14, 15^, suggesting a higher degree of accessibility to that site by alpha-mannosidases in the ER. The results of multi-microsecond simulations presented in this work indicate that a reduced degree of filling of the pocket left empty by the opening of the RBD by a smaller N-glycan at N243 leads progressively higher degree of instability of the wide-open conformation of the RBD. More specifically, while the N234-Man5 model appears competent in exposing the open RBD despite its higher dynamics relative to the N234-Man9 model, the N234-Man3 model leads to a dominant “more closed” conformation, see **Figure S.2**, where the paucimannose cannot form stable interactions within the pocket, interacting only with residues at the pocket’s gate or outside it, see **Figures 3** and **S.6**. The progressive destabilization upon reduction of the N-glycan size at N234 is in agreement with the results obtained for the N234A/N165A mutant^17^, designed to account for the complete removal of glycosylation at N234, which leads to 60% less binding to the ACE2 receptor as determined through biolayer interferometry assays^17^. It should be noted that the compact architecture of the S trimer does not allow for very large excursions in terms of lateral and especially of axial angles, nevertheless changes in these parameters in the same numerical range observed for the N234A/N165A mutant^17^ have been shown to correspond to dramatic difference in ACE2 binding.

In agreement with recent work that discusses the roles of the glycan at N343 in supporting the RBD intermediate dynamics between open and closed conformations^41^, and of the glycan at N165 in supporting the open RBD^17^, we observed that the stability of the N234-Man5 is significantly reduced when all the N-glycans in the shield are reduced to Man5 (All-Man5 model). In agreement with recent work^41^, this destabilization is due primarily to the lack of interactions that the shorter Man5 at N343 can make with the open RBD, and in particular with residues in the disordered loop within the RBM (400 to 508), in addition to the Man5 at N234 allowing the RBD to be relatively unhinged. We have shown that this destabilization is partially recovered when the N234 is occupied by a larger oligomannose, such as Man9. However, within this framework, the RBD adopts a more closed conformation, where the Man5 at both N165 and N343 can interact with the open RBD and with each other, see **Figure S.5**.

The introduction of an additional glycosylation site at N370, which SARS-CoV-2 S has lost due to the T372A mutation in the Wuhan-Hu-1 and derived strains, illustrates the importance of an effective filling of the cavity left empty by the opening of the RBD. Indeed, the complex FA2G2 glycan at N370 can easily access the pocket in addition to Man9 at N234, which in this specific case can only partially fill it due to steric hindrance. We have shown that the presence of an N370 glycan contributes effectively to the stability of the wide-open RBD state, which nevertheless needs to be achieved starting from a closed S conformation^41^. To this end, we find that the N370 N-glycan on the SARS-CoV-2 S throughout all our simulations occupies a specific cleft on the surface of the closed RBDs. This binding mode is stabilized by a network of hydrogen bonds and hydrophobic interactions between the protein and the monosaccharides of the FA2G2 N370 glycan (1-6) arms, see **Figure 4** and **S.7**. Because the N370 glycan involved in this interaction originates from the adjacent closed RBD, its binding results in tying the closed RBDs together, likely hindering opening in the first place. This is in agreement with recent work^50^ suggesting that the introduction of a N370 sequon in SARS-CoV-2 S negatively affects binding to human ACE2, contributing to increased replication of SARS-CoV-2 S in human cells relative to its putative ancestral variant. Furthermore, it is also interesting to note that cryo-EM S structures from the bat RatG13 and pangolin CoV variants, both carrying the N370 sequon, have been only solved in their closed states^25, 33, 34^, possibly also suggesting opening is less favoured in these S glycoproteins relative to the SARS-CoV-2 S.

Ultimately, the tight binding we observed between the N370 N-glycan and the surface of the SARS-CoV-2 S closed RBD surface is not only interesting in terms of the implications for higher ACE2-binding activity through loss of the corresponding sequon suggests, but also indicates the presence of a glycan binding site in that cleft, which is occupied in CoV variants that retain that sequon. In this context, recent work^35^ provided evidence that heparan sulfate (HS) binds the SARS-CoV-2 S RBD in a ternary complex with ACE2, as an essential interaction for cell infection, meanwhile unfractionated heparin, non-anti-coagulant heparin, heparin lyases, and lung HS are found to potently block SARS-CoV-2 S binding to ACE2 and infection. Notably, the same study provides evidence that SARS-CoV S binding to heparin-BSA is significantly reduced, yet not completely negated^35^. Furthermore, recent work has also shown evidence that the RBD of SARS-CoV-2 S specifically binds sialogangliosides, such as GM1 and GM2, with the same affinity observed for glycosaminoglycans (GAGs), as well as blood group antigens with a lower affinity^39^. Our results suggest that in SARS-CoV2 S a glycan-binding cleft on the RBD surface is available and broadly accessible to be occupied by GAGs, sialogangliosides as well as blood group antigens, provided that they fit the structural and electronic constraints that the site imposes. Meanwhile, in SARS-CoV S binding of these species may be disfavoured, because the cleft is occupied by the N370 glycan from the adjacent protomer, further stabilizing the closed conformation. Furthermore, because of the high density of GAGs and sialogangliosides displayed on the surface of mammalian cells, we can speculate that this recently acquired topological change of the glycan shield may be advantageous for the virus towards cell surface localization and increased affinity to ACE2, where these glycans act as co-receptors^35,39^. Further investigation on these topics is underway.

Analysis of reconstructed SARS S ancestral sequences indicates that while the N370 sequon was recently lost in SARS-CoV-2 S, this sequon was only quite recently acquired within the phylogeny. However, proximal N-glycosylation sequons for example at position 364 (D364-YS in CoV2) have been gained and lost alternatively, see **Figure 4 panel d**. Based on our results, it is reasonable to think that glycosylation within this S topological region may have been evolutionarily conserved, because of its role in effectively stabilizing the open RBD, despite the higher energetic cost involved in the transition from the RBD down-to-up state. In SARS-CoV-2, glycosylation at N234, which according to our analysis, shown in **Figure 4 panel d**, also appeared recently, would then functionally take the role of a glycan at N370. This evolutionary flexibility in the precise positions of glycosylation sites would make the presence of any specific glycan dispensable, with the consequent advantage of ensuring easier RBD opening reaction, and thus a more active S.

## Conclusions

In this work we have used multi-microsecond MD simulations to determine the effect of changes in the nature and topology of the SARS-CoV-2 S N-glycosylation at sites known to be involved in its function. Our results indicate that reducing the size of the N-glycans at N234 led to the instability of the “wide-open” RBD conformation, with a consequent increase in RBD dynamics and a progressive stabilization of conformations favouring the closed protomer. Additionally, the structure of the N-glycans at N165 and N343 also affects the stability of the open RBD with shorter structures unable to effectively interact with the RBD disordered loop within the RBM. This effect is especially dramatic when a shorter N-glycan, such as Man5, is also present at N234. To account for changes in the glycan shield topology, we explored the effect of re-introducing N-glycosylation to a recently lost sequon at N370. Our results indicate that while the N-glycan at N370 is highly effective in stabilizing the open RBD in conjunction with the N-glycan at N234, it tightly binds a specific cleft on the surface of the closed RBD, tying the closed protomers together and likely increasing the energetic cost of the RBD opening. Because the architecture of this RBD cleft is particularly able to bind multiple monosaccharides through a network of hydrogen bonds and dispersion interactions, we suggest that in SARS-CoV-2 it can be occupied by other diverse glycan structures, such as glycosaminoglycans, proteoglycans or sialylated species, which have also been shown to bind S. This flexibility in glycan-binding preference would provide an additional advantage in terms of increasing localization at the host cell surface. Finally, comparative analysis of reconstructed SARS ancestral sequences suggests that specific changes in the glycan shield topology at and around N370, in conjunction with the gain of N-glycosylation at N234, may have contributed to an increase in S activity, and thus of the infectivity of the SARS-CoV-2 relative to closely related coronaviruses.

## Supporting information

Movie S.1. Representation of the evolution of the N-glycosylation sequons (magenta) through the reconstructed SARS ancestral sequence on to the SARS-C

## Data Sharing

All MD trajectories are available OA in PDB format on https://github.com/CFogarty-2275/Fine_Tuning_The_Spike.

## Conflict of Interest

There are no conflicts to declare.

## Acknowledgements

The authors would like to thank Dr Lorenzo Casalino in Prof Rommie Amaro’s lab (UCSD) for sharing the *tcl* script for the calculation of the axial and lateral angle displacement of the RBD and Prof Amaro for insightful discussions. The Partnership for Advanced Computing in Europe (PRACE) is gratefully acknowledged for generous allocation of computational resources on Marconi100 hosted by CINECA, under the PRACE COVID-19 Fast Track scheme. CAF acknowledges the Irish Research council for funding under the Government of Ireland Postgraduate Scholarship scheme. AMH acknowledges Maynooth University (MU) for funding under the John and Pat Hume Postgraduate Scholarship scheme. AS acknowledges the Science Foundation Ireland for funding under Grant number 18/CRT/6049. The opinions, findings and conclusions or recommendations expressed in this material are those of the authors and do not necessarily reflect the views of the Science Foundation Ireland.

## Notes

### Competing Interest Statement

The authors have declared no competing interest.

## References

1. Y. Watanabe, T. A. Bowden, I. A. Wilson and M. Crispin, Biochim Biophys Acta Gen Subj, 2019, 1863, 1480–1497.

2. F. Li, Annu Rev Virol, 2016, 3, 237–261.

3. R. K. Plemper, Curr Opin Virol, 2011, 1, 92–100.

4. M. A. Tortorici and D. Veesler, Adv Virus Res, 2019, 105, 93–116.

5. A. C. Walls, M. A. Tortorici, J. Snijder, X. Xiong, B. J. Bosch, F. A. Rey and D. Veesler, Proc Natl Acad Sci U S A, 2017, 114, 11157–11162.

6. D. Wrapp, N. Wang, K. S. Corbett, J. A. Goldsmith, C. L. Hsieh, O. Abiona, B. S. Graham and J. S. McLellan, Science, 2020, 367, 1260–1263.

7. A. C. Walls, Y. J. Park, M. A. Tortorici, A. Wall, A. T. McGuire and D. Veesler, Cell, 2020.

8. H. Yao, Y. Song, Y. Chen, N. Wu, J. Xu, C. Sun, J. Zhang, T. Weng, Z. Zhang, Z. Wu, L. Cheng, D. Shi, X. Lu, J. Lei, M. Crispin, Y. Shi, L. Li and S. Li, Cell, 2020, 183, 730–738.e713.

9. B. Turoňová, M. Sikora, C. Schürmann, W. Hagen, S. Welsch, F. Blanc, S. von Bülow, M. Gecht, K. Bagola, C. Hörner, G. van Zandbergen, J. Landry, N. Trevisan Doimo de Azevedo, S. Mosalaganti, A. Schwarz, R. Covino, M. Mühlebach, G. Hummer, J. Krijnse Locker and M. Beck, Science, 2020, eabd5223.

10. Y. Watanabe, J. D. Allen, D. Wrapp, J. S. McLellan and M. Crispin, Science, 2020.

11. A. Shajahan, N. T. Supekar, A. S. Gleinich and P. Azadi, Glycobiology, 2020.

12. M. Sanda, L. Morrison and R. Goldman, Anal Chem, 2021, 93, 2003–2009.

13. P. Zhao, J. L. Praissman, O. C. Grant, Y. Cai, T. Xiao, K. E. Rosenbalm, K. Aoki, B. P. Kellman, R. Bridger, D. H. Barouch, M. A. Brindley, N. E. Lewis, M. Tiemeyer, B. Chen, R. J. Woods and L. Wells, Cell Host Microbe, 2020, 28, 586–601.e586.

14. Y. Watanabe, L. Mendonça, E. R. Allen, A. Howe, M. Lee, J. D. Allen, H. Chawla, D. Pulido, F. Donnellan, H. Davies, M. Ulaszewska, S. Belij-Rammerstorfer, S. Morris, A. S. Krebs, W. Dejnirattisai, J. Mongkolsapaya, P. Supasa, G. R. Screaton, C. M. Green, T. Lambe, P. Zhang, S. C. Gilbert and M. Crispin, ACS Cent Sci, 2021, 7, 594–602.

15. J. Brun, S. Vasiljevic, B. Gangadharan, M. Hensen, A. V Chandran, M. L. Hill, J. L. Kiappes, R. A. Dwek, D. S. Alonzi, W. B. Struwe and N. Zitzmann, ACS Cent Sci, 2021, 7, 586–593.

16. I. Bagdonaite, A. J. Thompson, X. Wang, M. Søgaard, C. Fougeroux, M. Frank, J. K. Diedrich, J. R. Yates, A. Salanti, S. Y. Vakhrushev, J. C. Paulson and H. H. Wandall, Viruses, 2021, 13.

17. L. Casalino, Z. Gaieb, J. A. Goldsmith, C. K. Hjorth, A. C. Dommer, A. M. Harbison, C. A. Fogarty, E. P. Barros, B. C. Taylor, J. S. McLellan, E. Fadda and R. E. Amaro, ACS Cent Sci, 2020, 6, 1722–1734.

18. T. Sztain, S. H. Ahn, A. T. Bogetti, L. Casalino, J. A. Goldsmith, E. Seitz, R. S. McCool, F. L. Kearns, F. Acosta-Reyes, S. Maji, G. Mashayekhi, J. A. McCammon, A. Ourmazd, J. Frank, J. S. McLellan, L. T. Chong and R. E. Amaro, Nat Chem, 2021.

19. J. Shang, Y. Wan, C. Luo, G. Ye, Q. Geng, A. Auerbach and F. Li, Proc Natl Acad Sci U S A, 2020, 117, 11727–11734.

20. M. Letko, A. Marzi and V. Munster, Nat Microbiol, 2020, 5, 562–569.

21. O. C. Grant, D. Montgomery, K. Ito and R. J. Woods, Sci Rep, 2020, 10, 14991.

22. M. I. Zimmerman, J. R. Porter, M. D. Ward, S. Singh, N. Vithani, A. Meller, U. L. Mallimadugula, C. E. Kuhn, J. H. Borowsky, R. P. Wiewiora, M. F. D. Hurley, A. M. Harbison, C. A. Fogarty, J. E. Coffland, E. Fadda, V. A. Voelz, J. D. Chodera and G. R. Bowman, Nat Chem, 2021.

23. Y. K. Choi, Y. Cao, M. Frank, H. Woo, S. J. Park, M. S. Yeom, T. I. Croll, C. Seok and W. Im, J Chem Theory Comput, 2021, 17, 2479–2487.

24. B. Coutard, C. Valle, X. de Lamballerie, B. Canard, N. G. Seidah and E. Decroly, Antiviral Res, 2020, 176, 104742.

25. A. G. Wrobel, D. J. Benton, P. Xu, C. Roustan, S. R. Martin, P. B. Rosenthal, J. J. Skehel and S. J. Gamblin, Nat Struct Mol Biol, 2020, 27, 763–767.

26. S. Neelamegham, K. Aoki-Kinoshita, E. Bolton, M. Frank, F. Lisacek, T. Lütteke, N. O’Boyle, N. H. Packer, P. Stanley, P. Toukach, A. Varki, R. J. Woods and S. D. Group, Glycobiology, 2019, 29, 620–624.

27. K. Cheng, Y. Zhou and S. Neelamegham, Glycobiology, 2017, 27, 200–205.

28. W. Humphrey, A. Dalke and K. Schulten, J Mol Graph, 1996, 14, 33-38, 27–38.

29. S. J. Williams and E. D. Goddard-Borger, Biochem Soc Trans, 2020, 48, 1287–1295.

30. J. Brun, S. Vasiljevic, B. Gangadharan, M. Hensen, A. Chandran, M. Hill, J. Kiappes, R. Dwek, D. Alonzi, W. Struwe and N. Zitzmann, bioRxiv, 2020.

31. W. T. Harvey, A. M. Carabelli, B. Jackson, R. K. Gupta, E. C. Thomson, E. M. Harrison, C. Ludden, R. Reeve, A. Rambaut, S. J. Peacock, D. L. Robertson and C.-G.U.C.-U. Consortium, Nat Rev Microbiol, 2021, 19, 409–424.

32. Y. Watanabe, Z. Berndsen, J. Raghwani, G. Seabright, J. Allen, J. McLellan, I. Wilson, T. Bowden, A. Ward and M. Crispin, bioRxiv, 2020.

33. A. G. Wrobel, D. J. Benton, P. Xu, L. J. Calder, A. Borg, C. Roustan, S. R. Martin, P. B. Rosenthal, J. J. Skehel and S. J. Gamblin, Nat Commun, 2021, 12, 837.

34. S. Zhang, S. Qiao, J. Yu, J. Zeng, S. Shan, L. Tian, J. Lan, L. Zhang and X. Wang, Nat Commun, 2021, 12, 1607.

35. T. M. Clausen, D. R. Sandoval, C. B. Spliid, J. Pihl, H. R. Perrett, C. D. Painter, A. Narayanan, S. A. Majowicz, E. M. Kwong, R. N. McVicar, B. E. Thacker, C. A. Glass, Z. Yang, J. L. Torres, G. J. Golden, P. L. Bartels, R. N. Porell, A. F. Garretson, L. Laubach, J. Feldman, X. Yin, Y. Pu, B. M. Hauser, T. M. Caradonna, B. P. Kellman, C. Martino, P. L. S. M. Gordts, S. K. Chanda, A. G. Schmidt, K. Godula, S. Leibel, J. Jose, K. D. Corbett, A. B. Ward, A. F. Carlin and J. D. Esko, Cell, 2020, 183, 1043–1057.e1015.

36. R. Tandon, J. S. Sharp, F. Zhang, V. H. Pomin, N. M. Ashpole, D. Mitra, M. G. McCandless, W. Jin, H. Liu, P. Sharma and R. J. Linhardt, J Virol, 2021, 95.

37. C. J. Mycroft-West, D. Su, I. Pagani, T. R. Rudd, S. Elli, N. S. Gandhi, S. E. Guimond, G. J. Miller, M. C. Z. Meneghetti, H. B. Nader, Y. Li, Q. M. Nunes, P. Procter, N. Mancini, M. Clementi, A. Bisio, N. R. Forsyth, V. Ferro, J. E. Turnbull, M. Guerrini, D. G. Fernig, E. Vicenzi, E. A. Yates, M. A. Lima and M. A. Skidmore, Thromb Haemost, 2020, 120, 1700–1715.

38. L. Liu, P. Chopra, X. Li, K. Bouwman, S. Tompkins, M. Wolfert, R. de Vries and G. Boons, ACS Cent Sci, 2021, 7, 1009–1018.

39. L. Nguyen, K. A. McCord, D. T. Bui, K. M. Bouwman, E. N. Kitova, M. Elaish, D. Kumawat, G. C. Daskhan, I. Tomris, L. Han, P. Chopra, T.-J. Yang, S. D. Willows, A. L. Mason, L. K. Mahal, T. L. Lowary, L. J. West, S.-T. D. Hsu, T. Hobman, S. M. Tompkins, G.-J. Boons, R. P. de Vries, M. S. Macauley and J. S. Klassen, Nat Chem Biol, 2021, 17.

40. A. N. Baker, S. J. Richards, C. S. Guy, T. R. Congdon, M. Hasan, A. J. Zwetsloot, A. Gallo, J. R. Lewandowski, P. J. Stansfeld, A. Straube, M. Walker, S. Chessa, G. Pergolizzi, S. Dedola, R. A. Field and M. I. Gibson, ACS Cent Sci, 2020, 6, 2046–2052.

41. T. Sztain, S. H. Ahn, A. T. Bogetti, L. Casalino, J. A. Goldsmith, R. S. McCool, F. L. Kearns, J. A. McCammon, J. S. McLellan, L. T. Chong and R. E. Amaro, bioRxiv, 2021.

42. K. M. Bouwman, I. Tomris, H. L. Turner, R. van der Woude, T. M. Shamorkina, G. P. Bosman, B. Rockx, S. Herfst, J. Snijder, B. L. Haagmans, A. B. Ward, G. J. Boons and R. P. de Vries, PLoS Pathog, 2021, 17, e1009282.

43. C. A. Fogarty and E. Fadda, J Phys Chem B, 2021.

44. C. O. Barnes, C. A. Jette, M. E. Abernathy, K. A. Dam, S. R. Esswein, H. B. Gristick, A. G. Malyutin, N. G. Sharaf, K. E. Huey-Tubman, Y. E. Lee, D. F. Robbiani, M. C. Nussenzweig, A. P. West and P. J. Bjorkman, Nature, 2020, 588, 682–687.

45. Y. Watanabe, L. Mendonça, E. R. Allen, A. Howe, M. Lee, J. D. Allen, H. Chawla, D. Pulido, F. Donnellan, H. Davies, M. Ulaszewska, S. Belij-Rammerstorfer, S. Morris, A. S. Krebs, W. Dejnirattisai, J. Mongkolsapaya, P. Supasa, G. R. Screaton, C. M. Green, T. Lambe, P. Zhang, S. C. Gilbert and M. Crispin, bioRxiv, 2021.

46. R. Parker, T. Partridge, C. Wormald, R. Kawahara, V. Stalls, M. Aggelakopoulou, J. Parker, R. Powell Doherty, Y. Ariosa Morejon, E. Lee, K. Saunders, B. F. Haynes, P. Acharya, M. Thaysen-Andersen, P. Borrow and N. Ternette, Cell Rep, 2021, 35, 109179.

47. Y. Watanabe, Z. T. Berndsen, J. Raghwani, G. E. Seabright, J. D. Allen, O. G. Pybus, J. S. McLellan, I. A. Wilson, T. A. Bowden, A. B. Ward and M. Crispin, Nat Commun, 2020, 11, 2688.

48. A. M. Harbison, L. P. Brosnan, K. Fenlon and E. Fadda, Glycobiology, 2019, 29, 94–103.

49. J. Pallesen, N. Wang, K. S. Corbett, D. Wrapp, R. N. Kirchdoerfer, H. L. Turner, C. A. Cottrell, M. M. Becker, L. Wang, W. Shi, W. P. Kong, E. L. Andres, A. N. Kettenbach, M. R. Denison, J. D. Chappell, B. S. Graham, A. B. Ward and J. S. McLellan, Proc Natl Acad Sci U S A, 2017, 114, E7348–E7357.

50. L. Kang, G. He, A. K. Sharp, X. Wang, A. M. Brown, P. Michalak and J. Weger-Lucarelli, Cell, 2021, 184, 4392–4400.e4394.

