## Supplementary figures and images for "Fine-tuning the Spike: Role of the nature and topology of the glycan shield in the structure and dynamics of the SARS-CoV-2 S"

### Movie S.1. Representation of the evolution of the N-glycosylation sequons (magenta) through the reconstructed SARS ancestral sequence on to the SARS-C

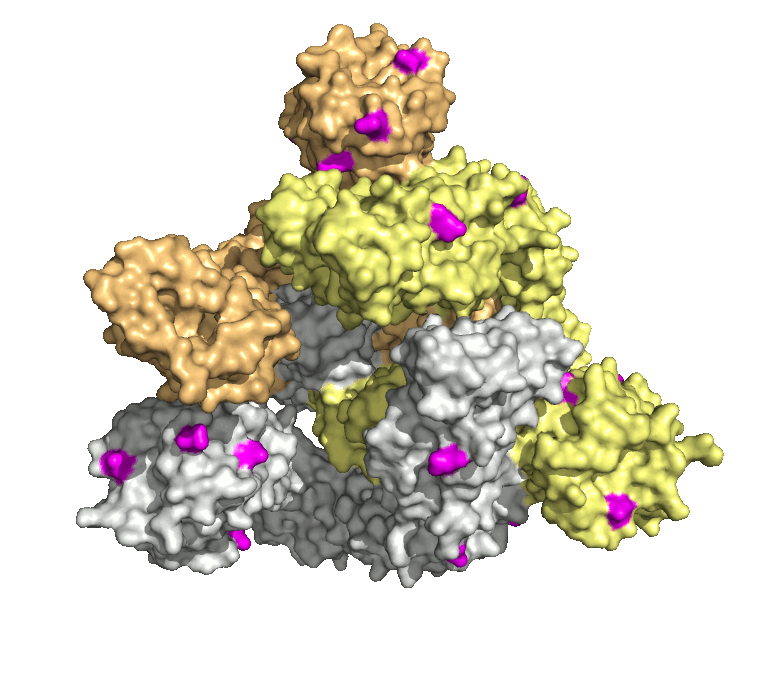
