## Supplementary Material for "Fine-tuning the Spike: Role of the nature and topology of the glycan shield in the structure and dynamics of the SARS-CoV-2 S"

**Computational Method**

**MD simulations: System set-up and protocol.** The SARS-CoV2 S models were generated by homology modelling using SWISS MODEL^1^ from the cryo-EM structure 6VYB (3.2 Å resolution) of the SARS-CoV-2 S 2P recombinant glycoprotein, with the reference sequence NCBI YP_009724390.1. To build our starting simulation model, the furin site was cleaved and the 2P mutation restored to the viral sequence. The loops not resolved in the cryo-EM structure were reconstructed using the SWISS MODEL structural libraries of backbone fragments, based on similar sequence structure. This resulted in a model with 54 N-glycosylation sites, 18 *per* protomer. The SARS-CoV2 S model, bearing the N370 sequon, was generated by introducing the A372T mutation. Complete glycoforms were reconstructed at the N-glycosylation sites, by aligning conformationally equilibrated N-glycan structures from our in-house *GlycoShape* library^2-4^ to the GlcNAc residues resolved in the cryo-EM structure, with slight adjustments of the torsion angles to resolve steric clashes with the surrounding protein, when necessary. Selection of the glycan at each N-glycosylation site was determined based primarily on work by Wantanabe *et al^5^*, with consideration to other studies available^6, 7^. O-glycosylation at T323 was chosen based on results from Shajahan *et al*^6^. The N-glycosylation at N370 in the SARS-CoV2 S mutant was chosen to be the same as reported for the SARS-CoV-1 S^8^. Summary of the specific glycosylation at each site is shown in **Table S.1** for all three protomers.

**Table S.1.** Site specific glycosylation chosen for the SARS-CoV-2 S glycoprotein models, with 18 N-glycan sites and 1 O-glycan site per protomer. Residue numbering (Resid) corresponds to the PDB 6VYB. The glycosylation sites where the glycosylation was changed between models are highlighted in red. N-glycans are represented by the Oxford nomenclature.

|  | N234-Man5 | N234-Man3 | N234-Man9 | N234-Man9* | N234-Man9**+N370 | N234-Man9* |
| --- | --- | --- | --- | --- | --- | --- |
| Resid | **Glycan Type** | | | | | |
| N61 | Man5 | | | | | |
| N74 | FA2G2 | | | | | |
| N122 | Man5 | | | | | |
| N149 | FA2G2 | | | | | |
| N165 | FA2G2 | | | A2B | A2B | Man5 |
| N234 | Man5 | Man3 | Man9 | | | |
| N282 | FA2G2 | | | | | |
| T323 | Neu5Ac-α3-Gal-β3-[Neu5Ac-α6]-GalNAc- | | | | | |
| N331 | FA2G2 | | | | | |
| N343 | FA2G2 | FA2G2 | FA2G2 | FA2B | FA2B | Man5 |
| N370 | *unoccupied* | | | | FA2G2 | *unoccupied* |
| N603 | Man5 | | | | | |
| N616 | A2G2 | | | | | |
| N657 | FA2G2 | | | | | |
| N709 | Man5 | | | | | |
| N717 | Man5 | | | | | |
| N801 | Man5 | | | | | |
| N1074 | Man5 | | | | | |
| N1098 | A2G2 | | | | | |
| N1134 | FA2G2 | | | | | |

* In this model the N-glycans at N165 and N234 are both Man5

** In this model the N-glycans at N165 and N234 are FA2B and A2B complex, with a bisecting GlcNAc, respectively.

In all MD simulations the protein and counterions (200 mM) were represented by the AMBER ff14SB^9^ parameter set, whereas the glycans were represented by the GLYCAM06j-1 version of the GLYCAM06 force field^10^. Water molecules were represented by the TIP3P model. All simulations were run with v18 of the AMBER software package^11^. The following running protocol was used for all MD simulations. The energy of the S ectodomains models was minimized in two steps of 50,000 cycles of the steepest descent algorithm each. During the first minimization all the heavy atoms were kept harmonically restrained using a potential weight of 5 kcal mol^−1^Å^2^, while the solvent, counterions and hydrogen atoms were left unrestrained. The minimization step was repeated with only the protein heavy atoms were kept restrained, while the glycans, solvent, counterions and hydrogens were left unrestrained. After energy minimization the system was equilibrated in the NVT ensemble with the same restraints scheme, where heating was performed in two stages over a total time of 1 ns, from 0 to 100 K (stage 1) and then 100 to 300 K (stage 2). During equilibration the SHAKE algorithm was used to constrain all bonds to hydrogen atoms. The Van der Waals interactions were truncated at 11 Å and Particle Mesh Ewald (PME) was used to treat long range electrostatics with B-spline interpolation of order 4. Langevin dynamics with collision frequency of 1.0 ps-1 was used to control temperature, which a pseudo-random variable seed to ensure there are no synchronization artefacts. Once the system was brought to 300 K an equilibration phase in the NPT ensemble of 1 ns was used to set the pressure to 1 atm. The pressure was held constant with isotropic pressure scaling and a pressure relaxation time of 2.0 ps. At this point all restraints on the protein heavy atoms were removed, allowing the system to evolve for 15 ns of conformational equilibration before production. At this stage different replicas for each model were generated starting from different velocities. The conformational equilibration phase for each replica was further extended to include the first 300 ns of production to allow the glycans shield to adapt to the protein architecture and vice-versa. In the analysis this 300 ns initial phase was discarded, see **Figure S.1**. The MD simulations were performed on PRACE ([www.prace-ri.eu](http://www.prace-ri.eu)) resources allocated on CINECA Marconi100, using 4 V100 GPUs per replica simulation, with a benchmark standard of approximately 25 ns/day. The total simulation times for each replica, including equilibration time, are shown in **Table 1** in the main manuscript. Average RMSD values and standard deviations are shown in **Table S.2**.

**Table S.2** Average RMSD values (Å) calculated for the RBD residues (330 to 530) in chain A, B and C, defined according to the original PDB used as a template to build the all SARS-CoV2 S models studied in this work. The RBDs in chains A and C are closed, while the RBD in chain B is open. Standard deviation values are shown in parenthesis.

| **Model System** | **RBD (A) *(closed)*** | **RBD (B) *(open)*** | **RBD (C) *(closed)*** |
| --- | --- | --- | --- |
| N234-Man5 (R1) | 5.8 (0.9) | 9.9 (1.1) | 4.6 (0.6) |
| N234-Man5 (R2) | 4.9 (0.6) | 9.8 (1.3) | 6.2 (0.8) |
| N234-Man5 (R3) | 6.5 (0.4) | 9.1 (0.8) | 4.1 (0.3) |
| N234-Man3 (R1) | 6.6 (0.9) | 12.2 (1.6) | 4.0 (0.7) |
| N234-Man3 (R2) | 6.2 (0.9) | 5.9 (1.3) | 4.0 (0.6) |
| N234-Man3 (R3) | 3.2 (0.8) | 10.7 (0.4) | 5.0 (0.2) |
| N234-Man9 (R1) | 4.6 (0.6) | 7.9 (0.7) | 4.1 (0.3) |
| N234-Man9 (R2) | 3.4 (0.4) | 6.9 (0.5) | 4.3 (0.2) |
| N234-Man9** N370 (R1) | 3.1 (0.4) | 12.8 (1.2) | 3.9 (0.7) |
| N234-Man9** N370 (R2) | 3.1 (0.6) | 4.0 (0.8) | 2.2 (0.4) |
| N234-Man9** | 2.6 (0.4) | 5.3 (0.6) | 3.3 (0.5) |
| N234-Man9** | 4.9 (0.4) | 8.8 (0.6) | 4.3 (0.4) |
| All Man5 (R1) | 5.4 (0.8) | 12.0 (1.0) | 3.6 (0.6) |
| All Man5 (R2) | 4.9 (0.4) | 11.5 (1.6) | 3.5 (0.3) |
| All Man5 (R3) | 3.9 (0.9) | 9.0 (1.7) | 4.2 (0.4) |
| N234-Man9* (R1) | 5.1 (0.6) | 9.5 (1.2) | 6.1 (0.7) |

* In this model the N-glycans at N165 and N234 are both Man5

** In this model the N-glycans at N165 and N234 are FA2B and A2B complex, with a bisecting GlcNAc, respectively.

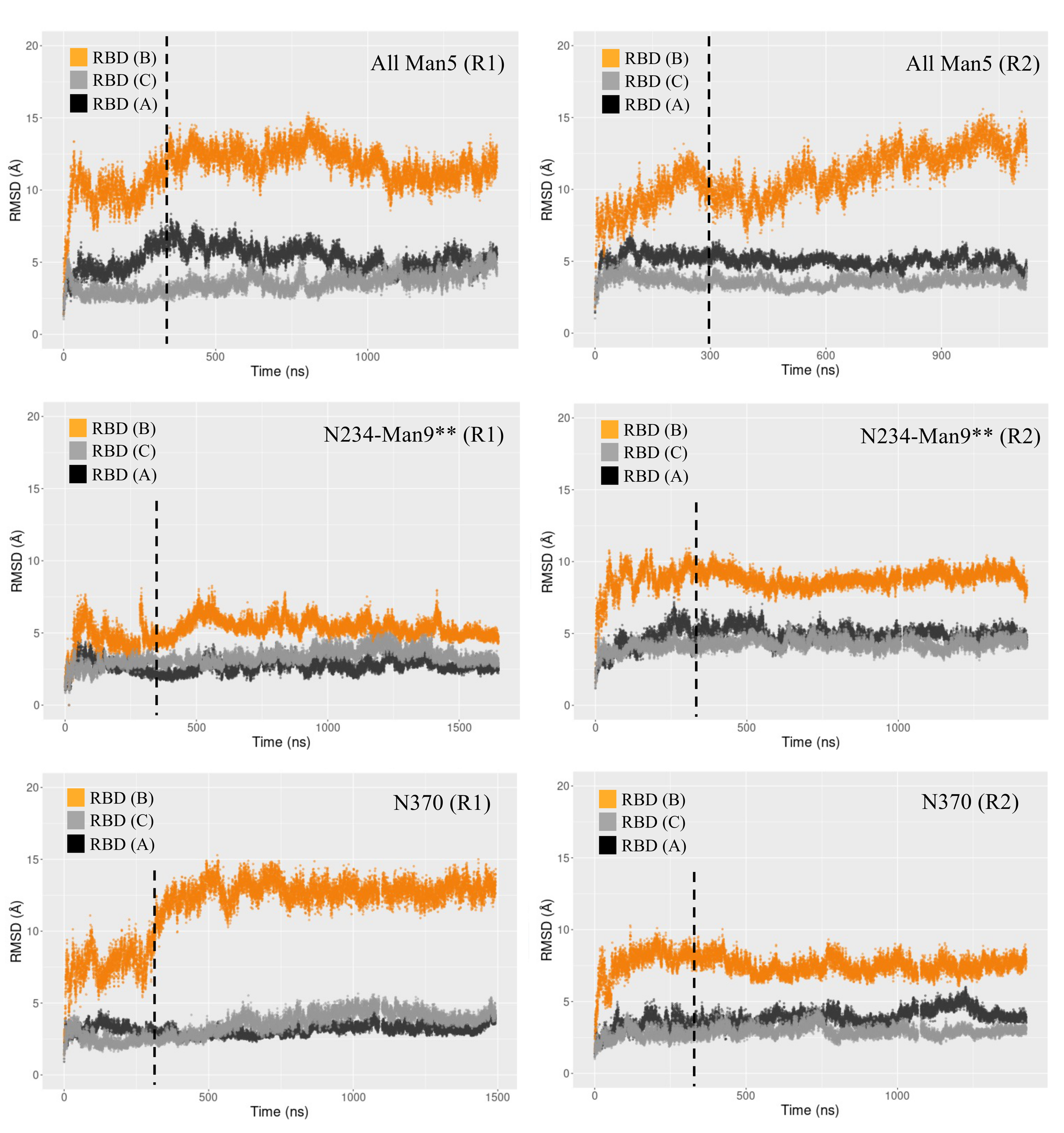

**Figure S.1** Time evolution of the three RBDs backbone root mean square deviation (RMSD) values for six simulations chosen as examples to illustrate our rationale for discarding the first 300 ns (dashed line) of the MD from the analysis, considering it as part of the conformational equilibration stage. Not in every case the 300 ns threshold is probably as necessary, see R1 and R2 N234 Man9** (see Table S.2 footnotes for nomenclature) and N370 (R2), which in these cases is further indication of the glycosylation supporting a “wide open” RBD corresponding to the cryo-EM PDB 6VYB used as a starting structure. Nevertheless, the same threshold was used for all simulations for consistency.

**Bioinformatics.** We obtained a list of SARS-CoV-2 S protein homologs from UniProt^11^ using a blastp search of the SARS-CoV-2 S protein sequence (UniProt Accession P0DTC2) against “Virus” proteins with an E-value threshold of 0.01, auto-selection of matrix, allowing gaps and with a maximum of 1000 hits. We further filtered this list of proteins to remove duplicates and only retain those with “spike” in the name. We aligned these proteins with Clustal Omega v1.2.2^12, 13^. A phylogenetic tree was created from the multiple sequence alignment using FastTree v2.1.10 without SSE3, and 1000 as the bootstrap parameter^14^. Ancestral sequence reconstruction was performed based on this multiple sequence alignment and the associated phylogenetic tree, using CodeML from the PAML v4.9e with WAG amino acid substitution matrix and molecular clock turning on^15^. A multiple sequence alignment of reconstructed sequences was parsed from the CodeML output. The locations of glycosylation sequons (Asn-Xaa-Ser/Thr, Xaa≠Pro) from the sequences before the alignment were mapped onto the aligned sequence positions including gaps in the aligned sequences. Using the script *spike_protein.tree_traversal.py,* we traversed the tree from the root to a specific extant sequence. This script utilized a list of extant proteins of interest, the multiple sequence alignment before ancestral sequence reconstruction, the multiple sequence alignment after ancestral sequence reconstruction, the phylogenetic tree including annotation of reconstructed nodes, and the generated dataset of N-linked sequons with their associated positions in the multiple sequence alignment after ancestral sequence reconstruction (including ancestral nodes). We recorded the status of each sequon from the ancestral sequence to each of its direct descendants, including whether the sequon had been lost, gained, or retained.

**Open RBD range of motions**

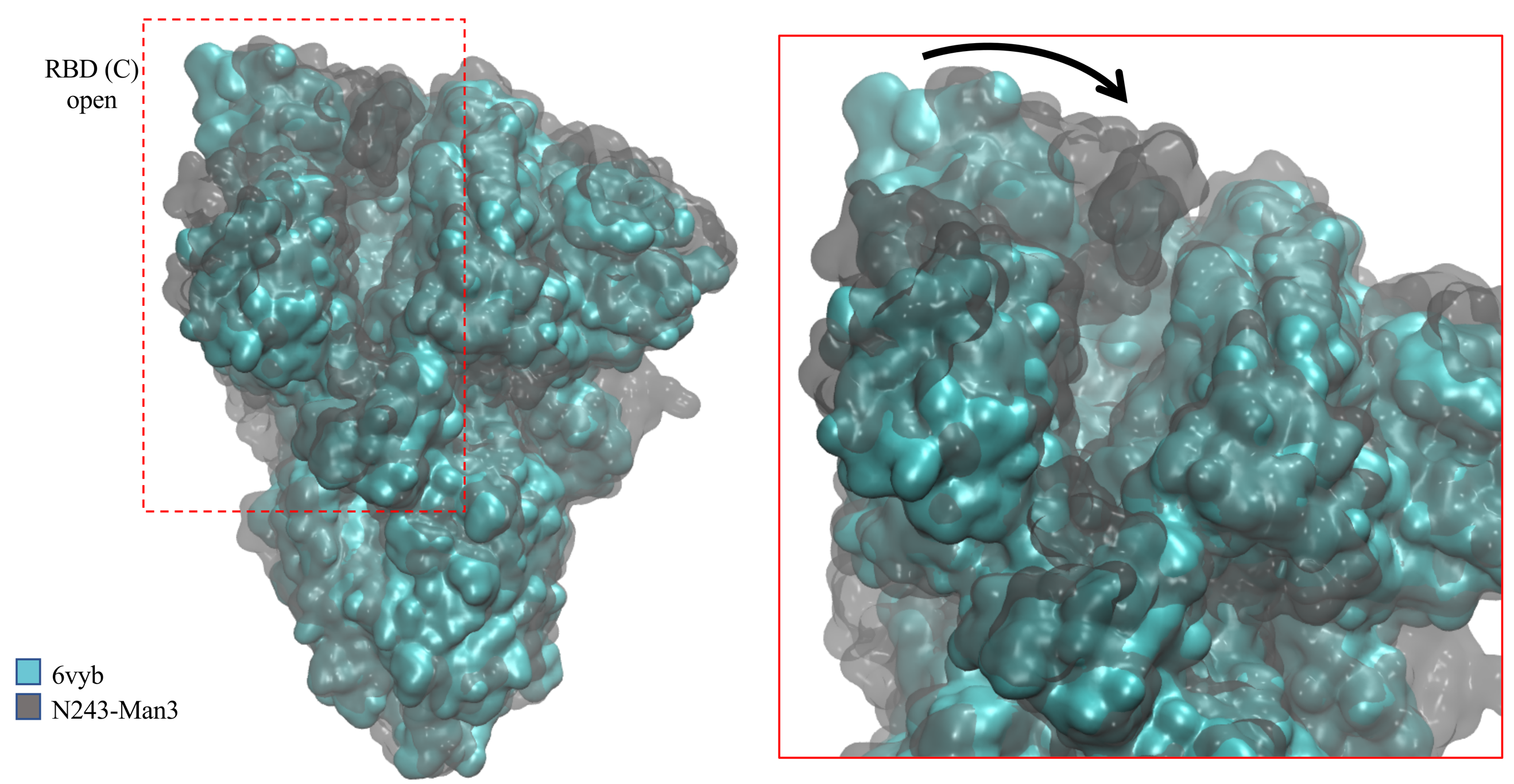

**Figure S.2.** Structural alignment (CA backbone) of the SARS-COV2 S cryo-EM structure PDBid 6vyb and a representative snapshot from the N234-Man3 trajectory (R1) to illustrate the wide range of motion of the open RBD (chain C in the PDB numbering) we observed in function of glycosylation. The proteins are represented by QuickSurf mode with VMD^16^, where 6vyb is rendered in cyan (opaque surface) and N234-Man3 in grey (glass3 surface). The insert on the right-hand side shows a close up of the section highlighted within the dashed rectangle on the figure, with an arrow as a visual guide connecting the two orientations. All glycans are omitted here for clarity.

**N234-Man5**

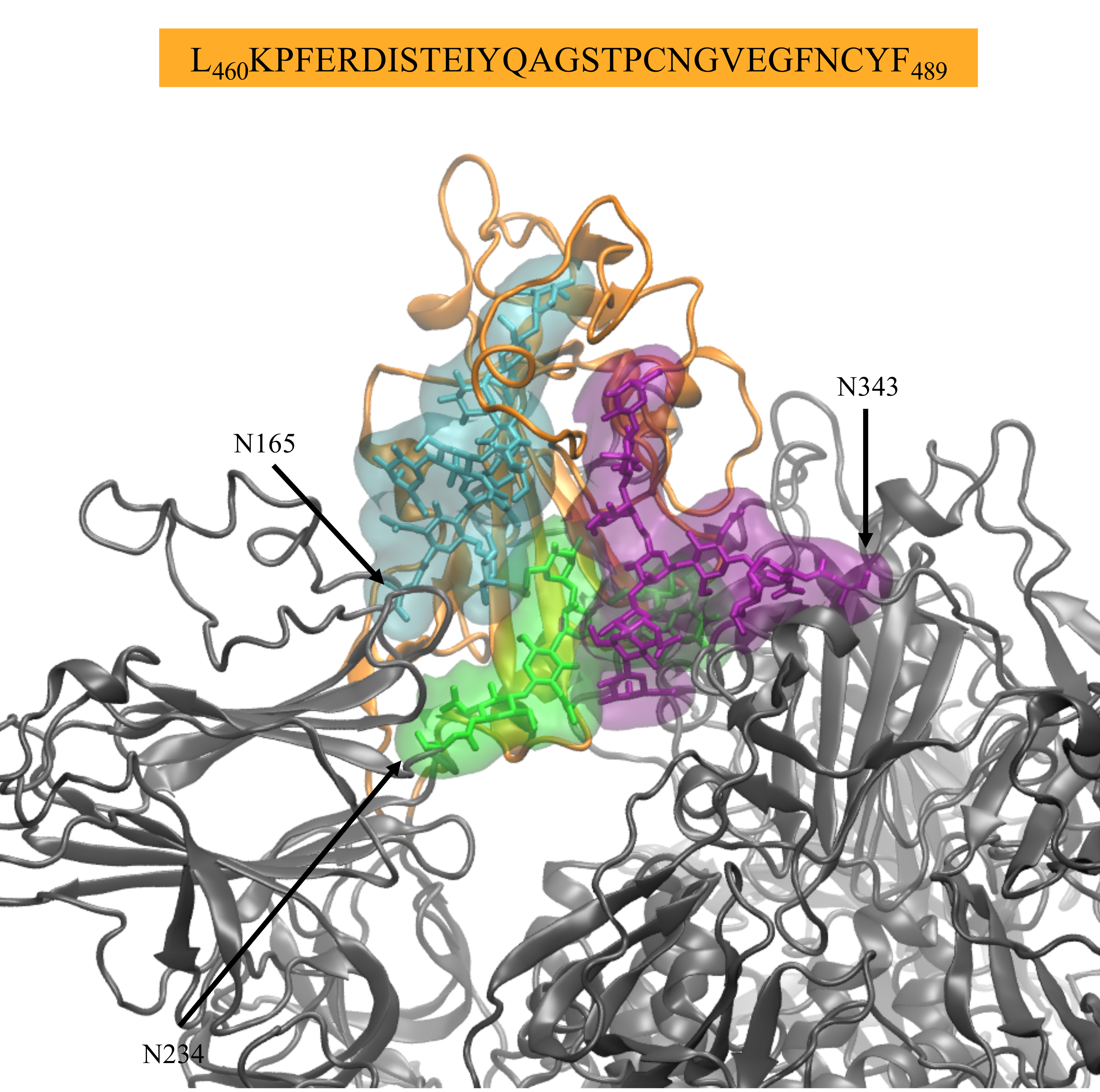

**Figure S.3** Close-ups on two representative snapshots of the N234-Man5 simulation R2 illustrating the orientation of the Man5 at N234 and corresponding interactions of the RBD (sequence indicated above) with the N-glycans at N165 and. N343. The Man5 at N234 is shown in green, while the FA2G2 at N165 and N343 are shown in cyan and purple, respectively. The open RBD (B) is shown in orange with a cartoon representation, while the rest of the protein is in grey. The glycans at all other sites are omitted for clarity. Molecular rendering with VMD^16^.

**All-Man5**

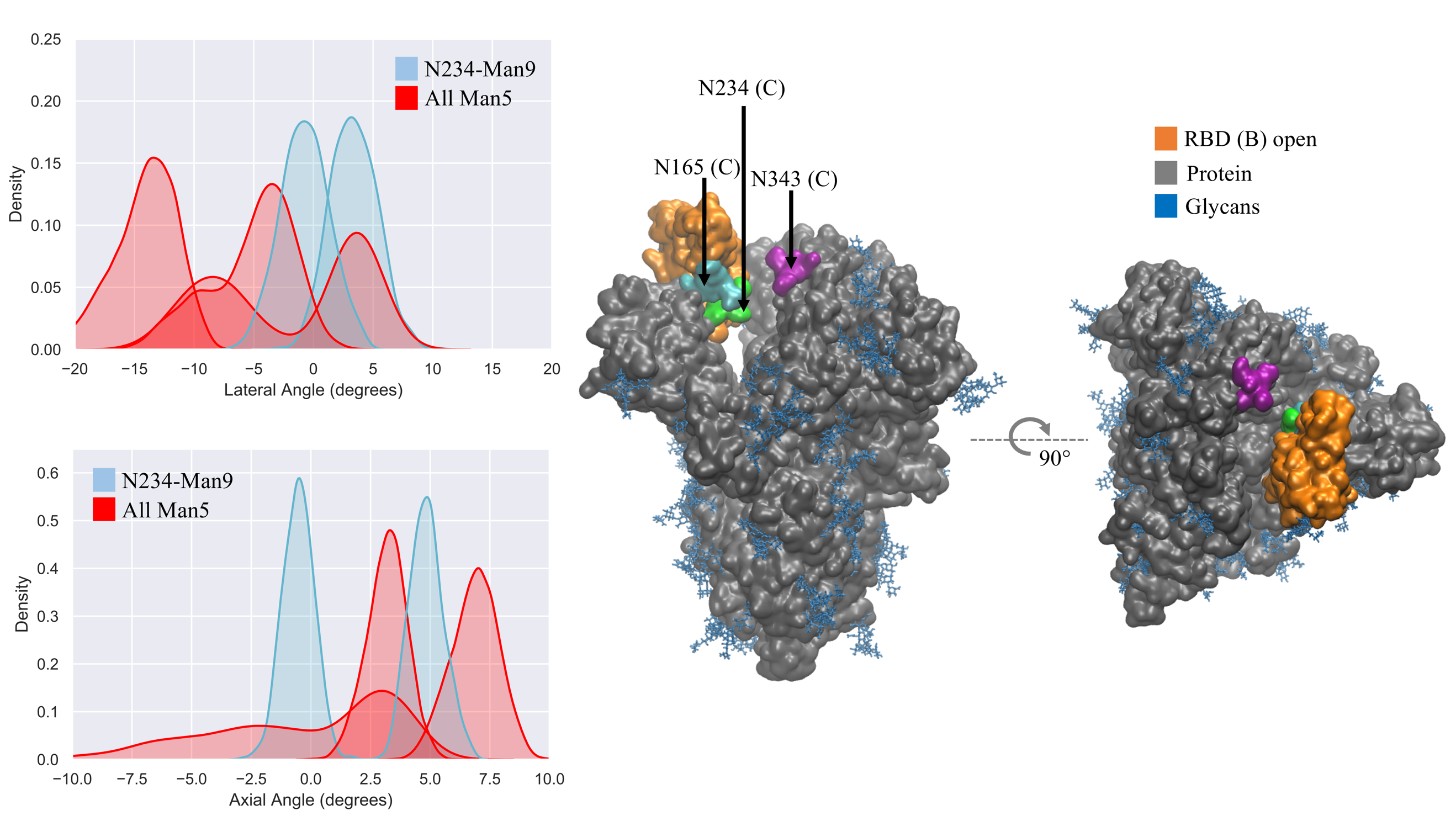

**Figure S.4. Left** KDE analysis of the lateral and axial angles distributions calculated through the uncorrelated MD trajectories obtained for All Man5 (red), replicas R1-3, and for N234-Man9 (cyan), replicas R1,2 for comparison. **Right** Close-ups on a representative snapshot of the All Man5 (R3) simulation from the side and top, respectively. Man5 at N234 is shown in green, while the Man5 at N165 and N343 are shown in cyan and purple, respectively. The solvent accessible surface of the open RBD is shown in orange, while the rest of the protein is shown in grey. All other glycans are shown in blue. Data analysis and graphs were done with seaborn ([www.seaborn.pydata.org](http://www.seaborn.pydata.org)) and molecular rendering with VMD^16^.

**N234-Man9* (N165/N343 Man5)**

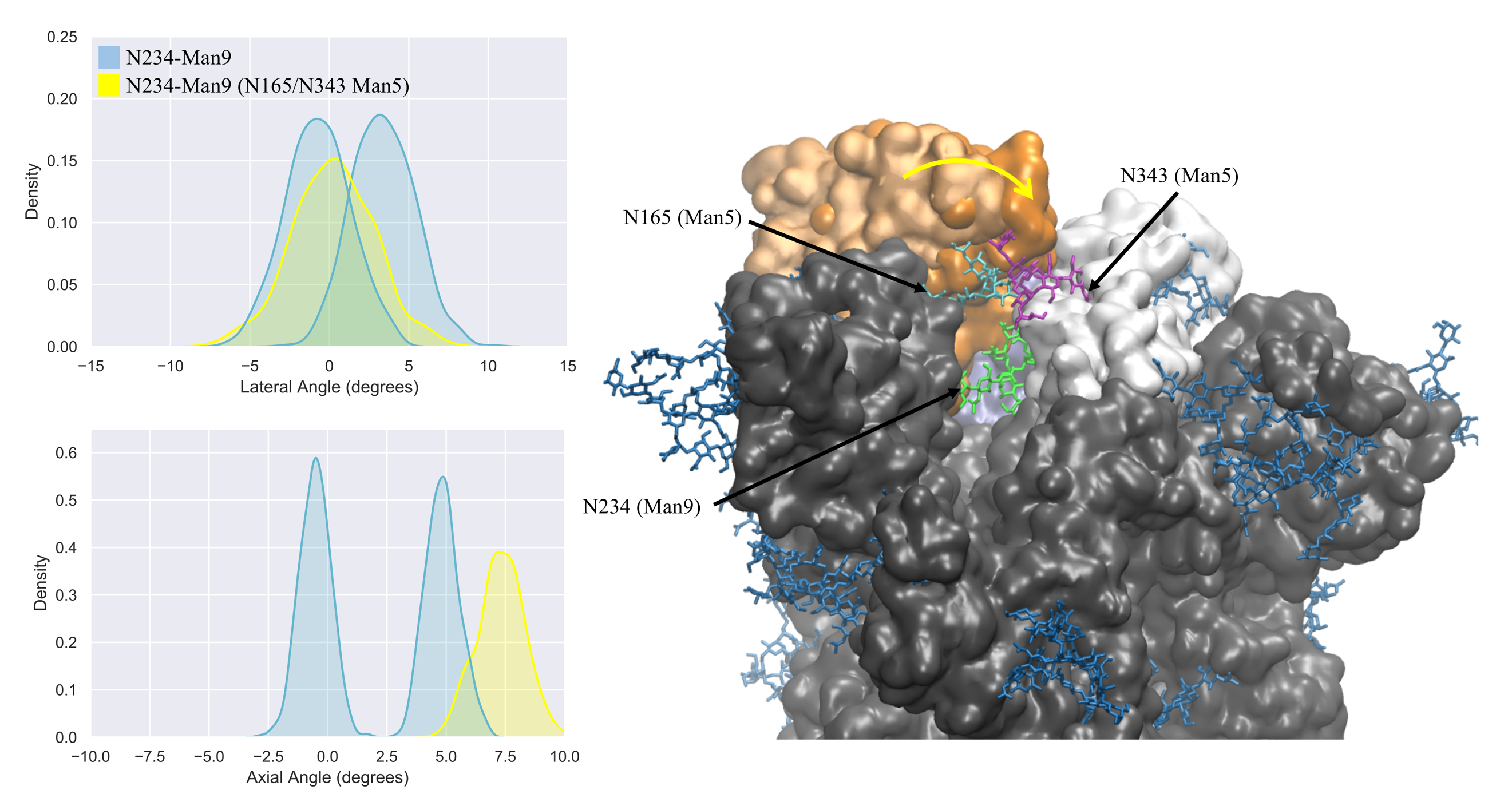

**Figure S.5. Left** KDE analysis of the lateral and axial angles distributions calculated through the uncorrelated MD trajectories obtained for N234-Man9 with Man5 at N165 and N343 (yellow), and for N234-Man9 (cyan), replicas R1,2 for comparison. **Right** Close-ups on a representative snapshot of the N234-Man9 with Man5 at N165 and N343 simulation from the side. Man9 at N234 is shown in green, while the Man5 at N165 and N343 are shown in cyan and purple, respectively. The solvent accessible surfaces of the open RBD (B) and closed RBD (C) are shown in orange and white, respectively. The displacement of the open RBD observed through the 2.1 μs trajectory is highlighted with a yellow arrow, where the initial RBD conformation is rendered in pale (brushed metal) orange. The rest of the protein is shown in grey and all other glycans are shown in blue. Data analysis and graphs were done with seaborn ([www.seaborn.pydata.org](http://www.seaborn.pydata.org)) and molecular rendering with VMD^16^.

**N234-Man3**

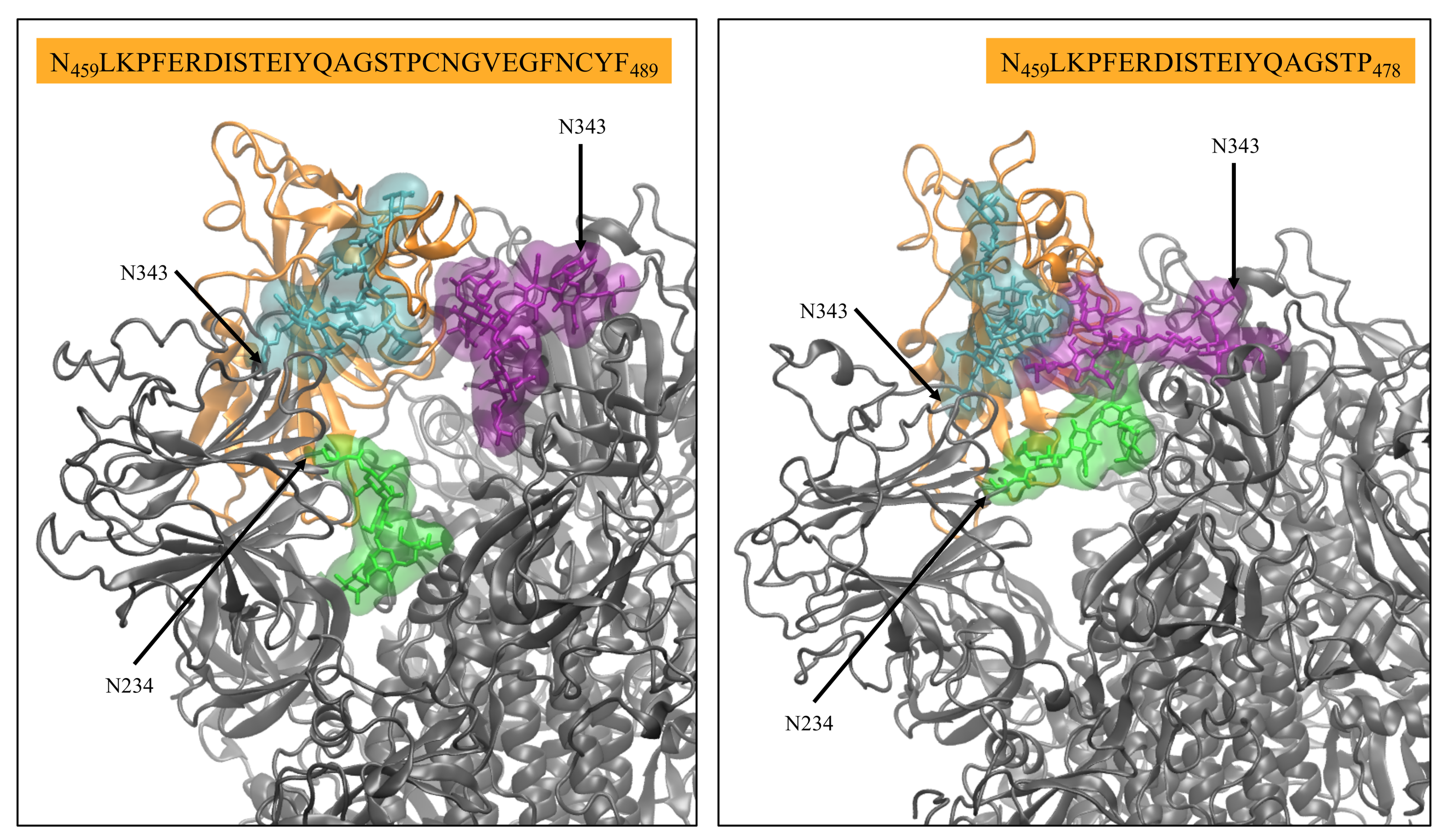

**Figure S.6.** Close-ups on two representative snapshots of the N234-Man3 simulations R1 (left) and R2 (right) illustrating different orientations of the Man3 at N234 and corresponding interactions of the RBD (sequence indicated above each panel) with the N-glycans at N165 and. N343. The Man3 at N234 is shown in green, while the FA2G2 at N165 and N343 are shown in cyan and purple, respectively. The open RBD (B) is shown in orange with a cartoon representation, while the rest of the protein is in grey. The glycans at all other sites are omitted for clarity. Molecular rendering with VMD^16^.

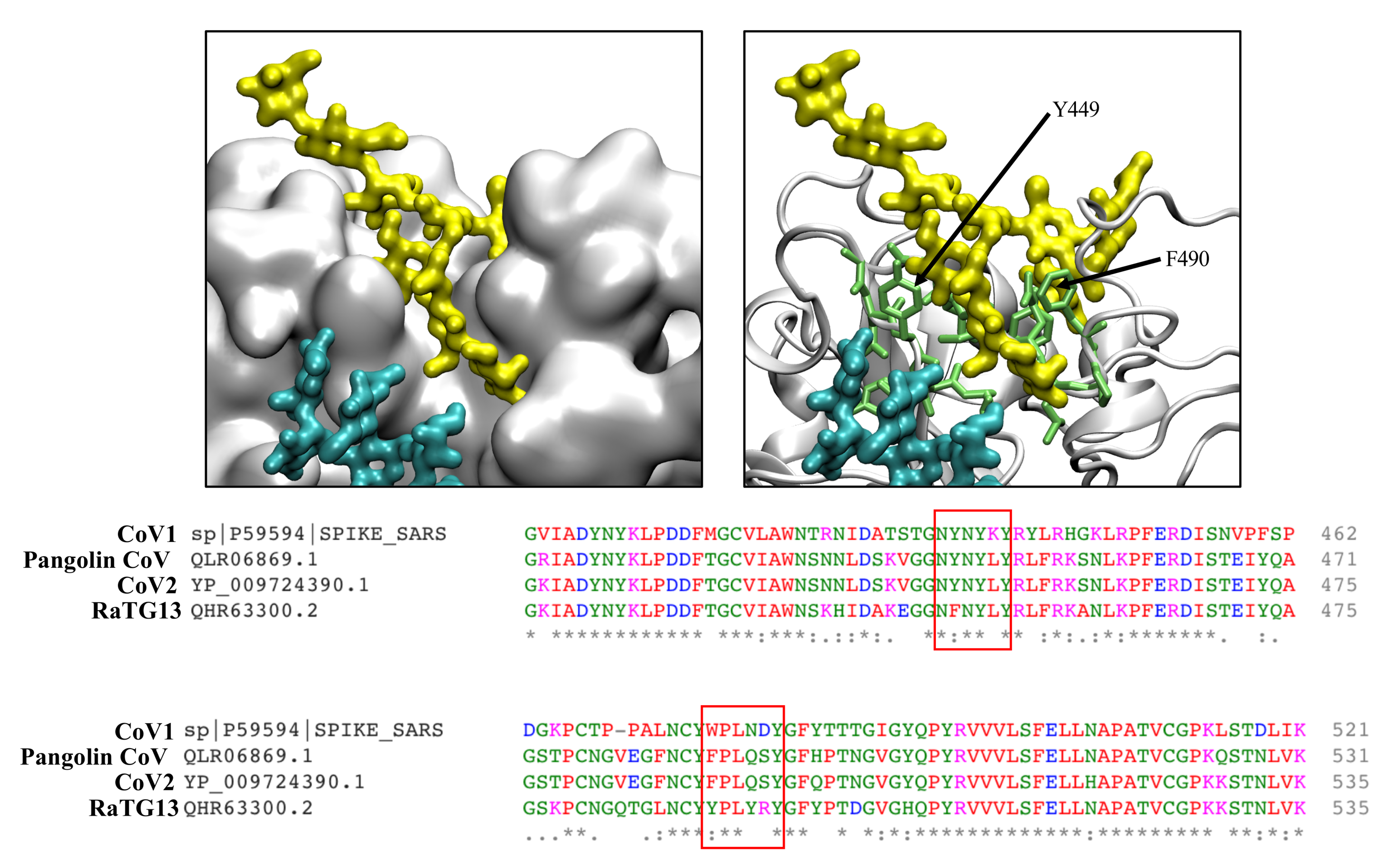

**Figure S.7** Close up of a snapshot of the MD simulation of the N234-Man9 glycosylated at N370 (R2). The FA2G2 N-glycan at N370 of chain A (yellow) is bound to the surface of the closed RBD (C) shown in as surface (left) in white. The N-glycan FA3G2 (bisecting GlcNAc) at N165 (C) is shown in cyan. On the top right image, the residues in contact with the N-glycan are shown in light green. The positions of Y449 and F490 are highlighted for ease of orientation. The sequence alignment of a section of the RBD in the CoV1, CoV2, pangolin CoV and of the bat RaTG13 S is shown at the bottom, where the residues lining the cleft, where the F2A2G2 (1-6) arm lies, are highlighted in light green. Rendering was done with VMD^16^ and sequence alignment with Clustal Omega^17^.

References

1.	Waterhouse, A.; Bertoni, M.; Bienert, S.; Studer, G.; Tauriello, G.; Gumienny, R.; Heer, F. T.; de Beer, T. A. P.; Rempfer, C.; Bordoli, L.; Lepore, R.; Schwede, T., SWISS-MODEL: homology modelling of protein structures and complexes. *Nucleic Acids Res* **2018,** *46* (W1), W296-W303.

2.	Fogarty, C. A.; Harbison, A. M.; Dugdale, A. R.; Fadda, E., How and why plants and human N-glycans are different: Insight from molecular dynamics into the "glycoblocks" architecture of complex carbohydrates. *Beilstein J Org Chem* **2020,** *16*, 2046-2056.

3.	Harbison, A. M.; Brosnan, L. P.; Fenlon, K.; Fadda, E., Sequence-to-structure dependence of isolated IgG Fc complex biantennary N-glycans: a molecular dynamics study. *Glycobiology* **2019,** *29* (1), 94-103.

4.	Fogarty, C. A.; Fadda, E., Oligomannose. *J Phys Chem B* **2021**.

5.	Watanabe, Y.; Allen, J. D.; Wrapp, D.; McLellan, J. S.; Crispin, M., Site-specific glycan analysis of the SARS-CoV-2 spike. *Science* **2020**.

6.	Shajahan, A.; Supekar, N. T.; Gleinich, A. S.; Azadi, P., Deducing the N- and O- glycosylation profile of the spike protein of novel coronavirus SARS-CoV-2. *Glycobiology* **2020**.

7.	Sanda, M.; Morrison, L.; Goldman, R., N- and O-Glycosylation of the SARS-CoV-2 Spike Protein. *Anal Chem* **2021,** *93* (4), 2003-2009.

8.	Watanabe, Y.; Berndsen, Z.; Raghwani, J.; Seabright, G.; Allen, J.; McLellan, J.; Wilson, I.; Bowden, T.; Ward, A.; Crispin, M., Vulnerabilities in coronavirus glycan shields despite extensive glycosylation. *bioRxiv* **2020**.

9.	Maier, J. A.; Martinez, C.; Kasavajhala, K.; Wickstrom, L.; Hauser, K. E.; Simmerling, C., ff14SB: Improving the Accuracy of Protein Side Chain and Backbone Parameters from ff99SB. *J Chem Theory Comput* **2015,** *11* (8), 3696-713.

10.	Kirschner, K. N.; Yongye, A. B.; Tschampel, S. M.; González-Outeiriño, J.; Daniels, C. R.; Foley, B. L.; Woods, R. J., GLYCAM06: a generalizable biomolecular force field. Carbohydrates. *J Comput Chem* **2008,** *29* (4), 622-55.

11.	Case, D.; Ben-Shalom, I.; Brozell, S.; Cerutti, D.; Cheatham III, T.; Cruzeiro, V.; Darden, T.; Duke, R.; Ghoreishi, D.; Gilson, M.; Gohlke, H.; Goetz, A.; Greene, D.; Harris, R.; Homeyer, N.; Izadi, S.; Kovalenko, A.; Kurtzman, T.; Lee, T.; LeGrand, S.; Li, P.; Lin, C.; Liu, J.; Luchko, T.; Luo, R.; Mermelstein, D.; Merz, K.; Miao, Y.; Monard, G.; Nguyen, C.; Nguyen, H.; Omelyan, I.; Onufriev, A.; Pan, F.; Qi, R.; Roe, D.; Roitberg, A.; Sagui, C.; Schott-Verdugo, S.; Shen, J.; Simmerling, C.; Smith, J.; Salomon-Ferrer, R.; Swails, J.; Walker, R.; Wang, J.; Wei, H.; Wolf, R.; Wu, X.; Xiao, L.; York, D.; Kollman, P. *AMBER 2018*, University of California, San Francisco, 2018.

12.	Bateman, A.; Martin, M.-J.; Orchard, S.; Magrane, M.; Agivetova, R.; Ahmad, S.; Alpi, E.; Bowler-Barnett, E. H.; Britto, R.; Bursteinas, B., UniProt: the universal protein knowledgebase in 2021. *Nucleic Acids Research* **2020**.

13.	Sievers, F.; Wilm, A.; Dineen, D.; Gibson, T. J.; Karplus, K.; Li, W.; Lopez, R.; McWilliam, H.; Remmert, M.; Söding, J., Fast, scalable generation of high‐quality protein multiple sequence alignments using Clustal Omega. *Molecular systems biology* **2011,** *7* (1), 539.

14.	Price, M. N.; Dehal, P. S.; Arkin, A. P., FastTree 2 – Approximately Maximum-Likelihood Trees for Large Alignments. *PLOS ONE* **2010,** *5* (3), e9490.

15.	Yang, Z., PAML 4: phylogenetic analysis by maximum likelihood. *Molecular biology and evolution* **2007,** *24* (8), 1586-1591.

16.	Humphrey, W.; Dalke, A.; Schulten, K., VMD: visual molecular dynamics. *J Mol Graph* **1996,** *14* (1), 33-8, 27-8.

17.	Sievers, F.; Wilm, A.; Dineen, D.; Gibson, T.; Karplus, K.; Li, W.; Lopez, R.; McWilliam, H.; Remmert, M.; Soding, J.; Thompson, J.; Higgins, D., Fast, scalable generation of high-quality protein multiple sequence alignments using Clustal Omega. *Molecular Systems Biology* **2011,** *7*.

   ^‡^ These authors equally contributed to the work presented in this manuscript
